# Hidden Structural States of Proteins Revealed by Conformer Selection with AlphaFold-NMR

**DOI:** 10.1101/2024.06.26.600902

**Authors:** Yuanpeng J. Huang, Theresa A. Ramelot, Laura E. Spaman, Naohiro Kobayashi, Gaetano T. Montelione

## Abstract

We introduce AlphaFold-NMR, a novel approach to NMR structure determination that reveals previously undetected protein conformational states. Unlike conventional NMR methods that rely on NOE-derived spatial restraints, AlphaFold-NMR combines AI-driven conformational sampling with Bayesian scoring of realistic protein models against NOESY and chemical shift data. This method uncovers alternative conformational states of the enzyme *Gaussia* luciferase, involving large-scale changes in the lid, binding pockets, and other surface cavities. It also identifies similar yet distinct conformational states of the human tumor suppressor Cyclin-Dependent Kinase 2-Associated Protein 1. These studies demonstrate the potential of AI-based modeling with enhanced sampling to generate diverse structural models followed by conformer selection and validation with experimental data as an alternative to traditional restraint-satisfaction protocols for protein NMR structure determination. The AlphaFold-NMR framework enables discovery of conformational heterogeneity and cryptic pockets that conventional NMR analysis methods do not distinguish, providing new insights into protein structure-function relationships.

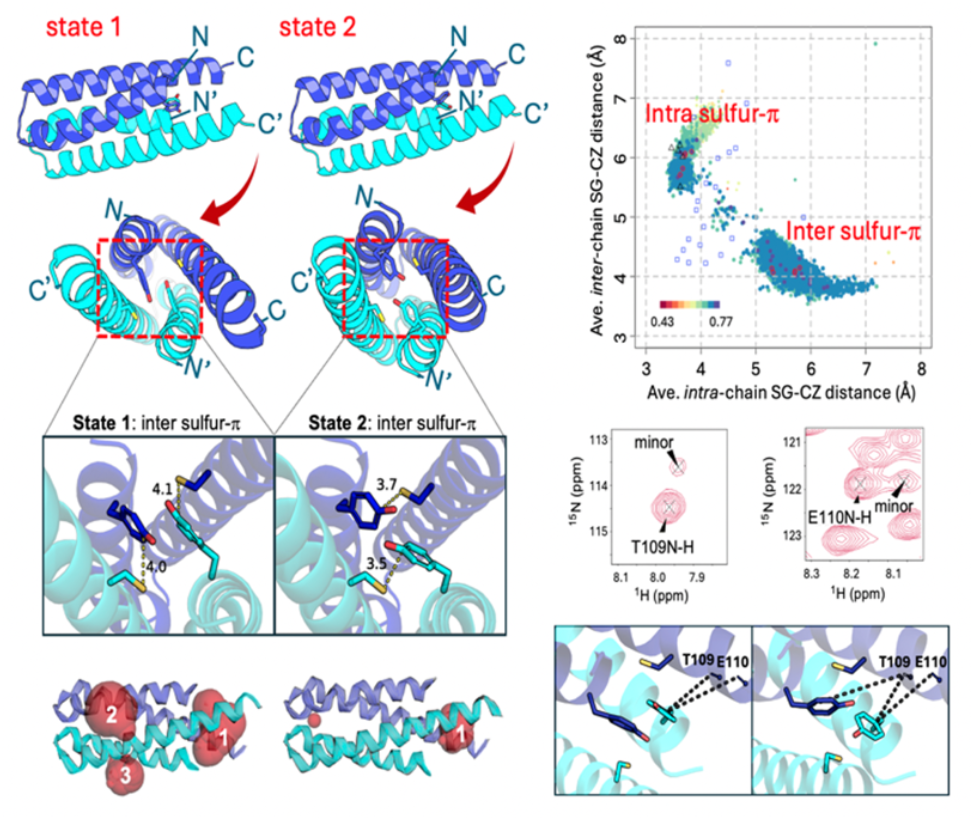

## Introduction

Recent advances in molecular modeling using artificial intelligence (AI), such as AlphaFold2 (AF2), have revolutionized protein structure determination. For example, AF2 has demonstrated high accuracy in modeling relatively rigid protein structures^1^. AF2 models are now widely used as phasing models in X-ray crystallography^2,3^ and to guide structural analysis in cryogenic electron microscopy (cryoEM), small angle X-ray scattering (SAXS), and NMR [for examples see refs ^4-8^]. While AF2 was not designed for studying protein dynamics, it has been adapted to predict flexible regions^1,9-13^, alternative conformations^14-22^, and populations of conformational states^23-25^ of proteins. However, the general problem of identifying biologically relevant structures from these computational ensembles remains challenging, particularly for dynamic proteins.

To address this challenge, we developed AlphaFold-NMR (AF-NMR), which combines AI-driven enhanced sampling with NMR-guided conformational selection. First, an enhanced sampling AF2 protocol is used to generate a wide range of physically-reasonable structural models. The generated models are clustered, scored against NMR data, and validated using NOESY Double Recall analysis. Compared to conventional NMR restraint-based modeling, AF-NMR overcomes inaccuracies that result from interpreting NOESY data as distance restraints^26,27^, particularly those arising from conformational averaging, and enables accurate determination of both single and multi-state protein structures. While AFsample^21^, a neural network dropout method, is used here for enhanced sampling; other comprehensive sampling strategies could also be incorporated into our protocol. This approach provides a robust framework for studying dynamic proteins and uncovering otherwise hidden conformational states of proteins.

We selected *Gaussia* luciferase (GLuc) for NMR-guided conformational selection due to its dynamic nature and likelihood of sampling multiple states in equilibrium^28,29^. GLuc, an 18.2 kDa monomeric enzyme from *Gaussia princeps*, catalyzes the oxidation of coelenterazine, producing a bright blue bioluminescent signal^30^. Previous NMR studies have highlighted its extensive conformational flexibility (PDB_ID: 7d2o^28,31^; PDB_ID: 9lfa^32^), and it was classified as a challenging target in CASP14 (target T1027) due to the absence of homologous structure templates for comparative modeling^29^. Notably, there are significant structural differences between the computed AF2 model and the experimental NMR structure, particularly in the positions of three helices^29^. This discrepancy is of interest because AF2 models of small, relatively rigid proteins often rival the accuracy of experimental NMR structures^4,6,7,29,33,34^, despite NMR structures being excluded from AF2 training^1^. However, for highly-flexible proteins like GLuc, these differences highlight AF2’s limitations in capturing the solution-dominant states^11,29,35-38^. Both GLuc models are supported by distinct subsets of NOEs, leading us to reason that NOESY data could be used for conformational selection. These characteristics make GLuc an excellent system for exploring NMR-guided AI-enhanced sampling methods for modeling multiple conformational states of proteins.

Using AF-NMR, we uncovered multiple conformational states of GLuc that collectively fit NOESY and chemical shift data better than the original restraint-generated NMR structure. A single set of chemical shifts indicates that these conformations are in fast -exchange on the NMR timescale. In one state, a loop between helices H5 and H6 adopts a closed conformation, forming a lid over a cryptic pocket proposed to be the enzyme’s active site^31,32^. In a second data-supported state, the lid is open, exposing the pocket, while subtle helical rearrangements further expand the pocket. A second surface pocket containing additional functionally-important residues is also observed. Comparing the AF-derived open and closed states to the NOESY data reveals distinct subsets of NOEs unique to each conformation, validating these two previously unrecognized structural states.

As a second example of AF-NMR revealing hidden structural states, we examined the human tumor suppressor Cyclin-Dependent Kinase 2-Associated Protein 1 (CDK2AP1). CDK2AP1, encoded by the *doc-1* (deleted in oral cancer 1) gene, is a specific inhibitor of CDK2^39^. Its loss or downregulation is associated with increased metastasis in various cancers, including liver and breast cancer ^40-42^. CDK2AP1 is a highly conserved protein that functions as both a cell cycle and epigenetic regulator^43^. It has multiple binding partners and forms a subunit of the nucleosome remodeling and histone deacetylation (NuRD) complex^44-46^, which regulates embryonic stem cell differentiation. The previously determined NMR structure of CDK2AP1 revealed a homodimeric four-helix bundle (residues 61-115) with intrinsically disordered 60-residue N-terminal regions^47^. Unlike GLuc, which exhibits large-scale structural heterogeneity, the flexibility of the CDK2AP1_61-115_ four-helical bundle is largely confined to its C-terminal helices, as indicated by ^15^N relaxation experiments^47^. Although Cys105 is considered essential for function and has been proposed to form an interchain disulfide bond^48^, the solution NMR structure indicates that these sulfur atoms are too far apart (4.4 - 10.2 Å) to form a covalent bond^47^. Using AF-NMR, we identified two conformational states of CDK2AP1_61-115_ in dynamic equilibrium, distinguished by interchain and intrachain Cys S– aromatic π interactions. These alternative structural states suggest a previously unrecognized role for Cys105: dynamically switching between interchain and intrachain sulfur–π interactions with Tyr63, and reveal previously hidden cryptic pockets on the surface of the CDK2AP1. These examples highlight the utility of AF-NMR for uncovering conformational states that fit to experimental data by *conformer selection*, offering a powerful alternative to conventional restraint-based protocols for protein NMR structure determination.

## Results

### Comparison of conventional NMR and standard AF2 models of GLuc using NOESY Double Recall

Discrepancies between AF2 models and experimental NMR structures for GLuc, as well as the identification of distinct NOEs supporting each model^29^, motivated its selection for this study. **Figure 1a** compares the conventional NMR structure (PDB ID 7d2o^28^) and models generated using AF2 Colabfold (see Methods). NOESY Double Recall analysis (**Figure 1b**) highlights regions of structural differences supported by NOESY peak data, with 543 distinct NOEs for the NMR model (blue dots) and 138 distinct NOEs for the AF2 models (orange dots). Key structural differences include the packing of N-terminal helix H1 inside the core (in NMR_7d2o_) versus helices H10/H11 packing against the core (in the standard AF2 model).

**Fig. 1.**
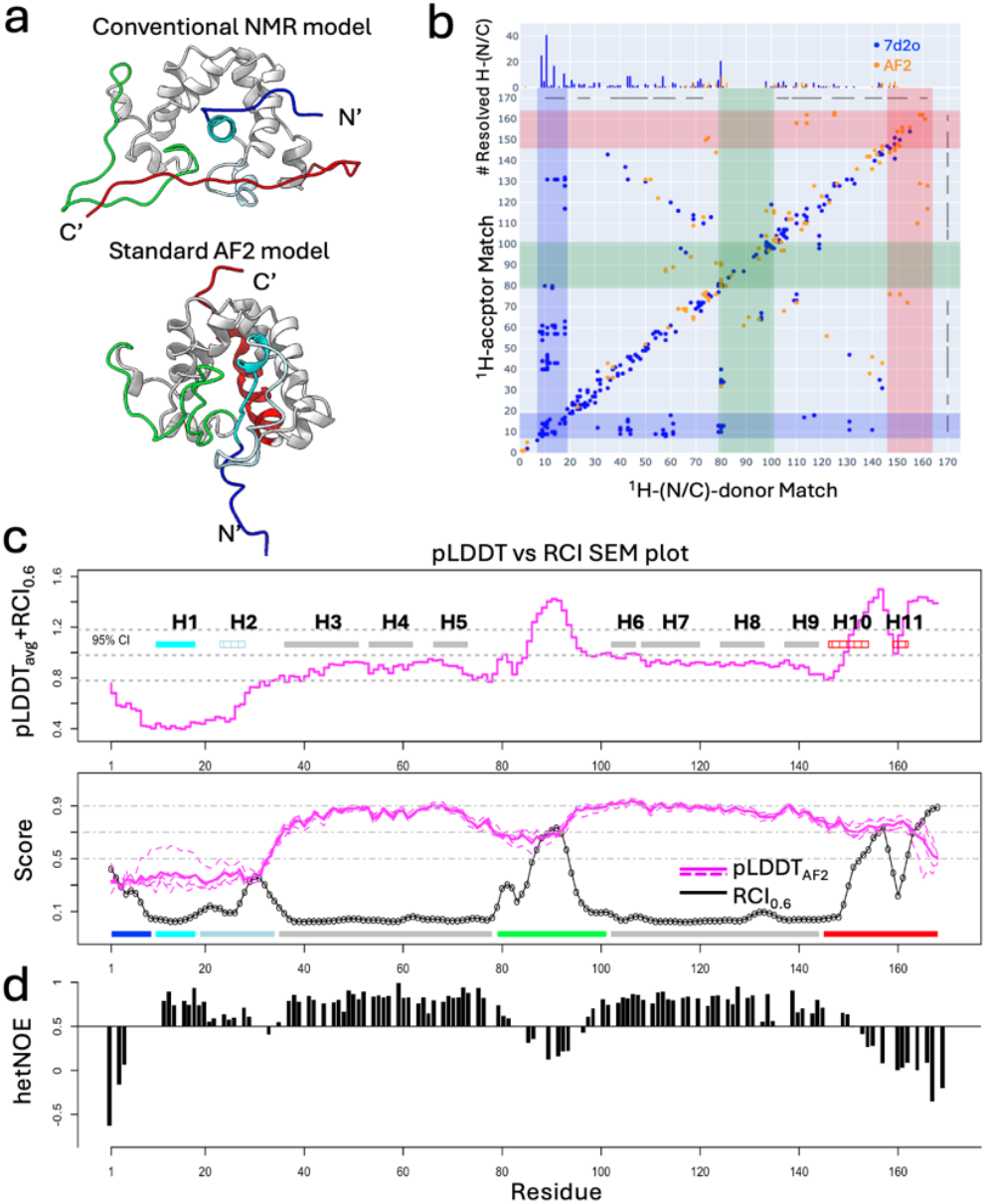
Comparison of conventional NMR and AF2 models of GLuc. **(a)** Ribbon diagrams of the NMR_7d2o_ structure (top) and standard AF2 model (bottom) with the structurally variable regions colored and conserved regions in gray. Color scheme: blue (N-terminal disordered region, residues 1-9), cyan (helix H1, residues 10-18), light blue (helix H2 and loop, residues 19-34), green (loop between H5 and H6, residues 79-101), red (C-terminal disordered region, residues 146-168), and gray (core α-helices H3 - H9). In the AF2 models, the C-terminal region (red) forms a kinked “broken helix” (residues 146-154 and 159-162). **(b)** Double Recall plot (see Methods) comparing NMR and AF2 models. Contact maps show inter-residue NOEs unique to the NMR model (blue), AF2 model (orange), or overlapped (brown). Blue and red stripes highlight contacts between the N-terminal helix H1 (residues 7-19) or C-terminal helices H10/H11 (residues 146-164) and core residues. **(c)** Comparison of pLDDT and RCI data. (Top) Comparison SEM plot (see Methods) between pLDDT_avg_ and RCI_0.6_ with helices labeled. The most flexible helices (H2, H10, and H11) are marked with stripes. The average and 95% confidence intervals (**Eqn. 3)** are shown as gray dashed lines. (Bottom) Per-residue pLDDT (dashed line for each model and solid line for average) and RCI_0.6_ scores. The residue segment color scheme of **Fig. 1a** is shown along the bottom. **(d)** Previously published per-residue backbone heteronuclear ^15^N-^1^H hetNOEs monitoring backbone dynamics^28^.

A more recent solution NMR structure of GLuc, PDB ID 9fla^32^, exhibits similar core architecture (**Supplementary Figure S1**). Both NMR_7d2o_ and AF2 structures share five disulfide bonds and a scaffold of seven helices, defined as “core helices”, although some significant modeling differences are reported. For comparisons, the NMR_7d2o_ structure was chosen over NMR_9fla_, as it has more complete and accessible NOESY and chemical shift data.

### Comparison of conventional NMR and standard AF2 models of GLuc using flexibility metrics

Experimental flexibility metrics, including the random coil backbone chemical shift index (RCI) and ^15^N-^1^H heteronuclear NOE (hetNOE), were compared with AF2 pLDDT scores, a measure of local model uncertainty (**Figure 1c,d**). While pLDDT scores generally correlate with regions of conformational flexibility^1,9-13,49^, this correlation is appreciated to be imperfect and fails to capture gradations in dynamics within highly flexible regions^13^. For GLuc, RCI and hetNOE profiles are strongly correlated (Spearman correlation coefficient, SCC: −0.75), while RCI and per-residue pLDDT values show weaker correlations (SCC: −0.56; **Figure 1c**).

The pLDDT vs RCI comparison plot (see Methods) illustrates structural regions of concordance and discrepancy, with good agreement in the rigid core (helices H3–H5 and H6–H9), where high pLDDT scores, low RCI values, and high hetNOE values are observed (**Figure 1c,d**). In contrast, flexible regions, such as the H5/H6 loop, have high RCI and low hetNOE values but relatively high pLDDT scores, suggesting an underrepresentation of flexibility in the AF2 models. Similarly, helix H1 has low RCI and high hetNOE values (indicating low flexibility) but exhibits low pLDDT scores in standard AF2 models. These discrepancies correspond to regions of significant structural differences between AF2 and NMR models (**Figure 1a**), suggesting the presence of multiple conformational states.

### AI-based modeling with enhanced sampling generates conformational diversity of GLuc

We tested whether AI-based modeling with enhanced sampling could produce alternative models that better fit the NMR NOESY and RCI data compared to standard AF2 models. This approach, summarized in **Supplementary Figure S2**, uses the AFsample method^21,50^, which generates diverse models by varying parameter settings. Since GLuc has a relatively shallow MSA (229 sequences), MSA-based methods such as AF_alt^14^ and AF_cluster^20^ failed to generate models with high NOESY recall scores and were not considered further. The resulting AFsample models were evaluated by TM score and NOESY recall scores (**Figure 2a**), which quantify the fraction of NOESY peaks explained by short interatomic distances. While most AFsample models had lower NOESY recall scores, some achieved scores as high as 0.88, surpassing the standard AF2 model (0.86) and approaching the conventional NMR models (0.89) that have been forced to fit as many NOESY-based restraints as possible.

**Fig. 2.**
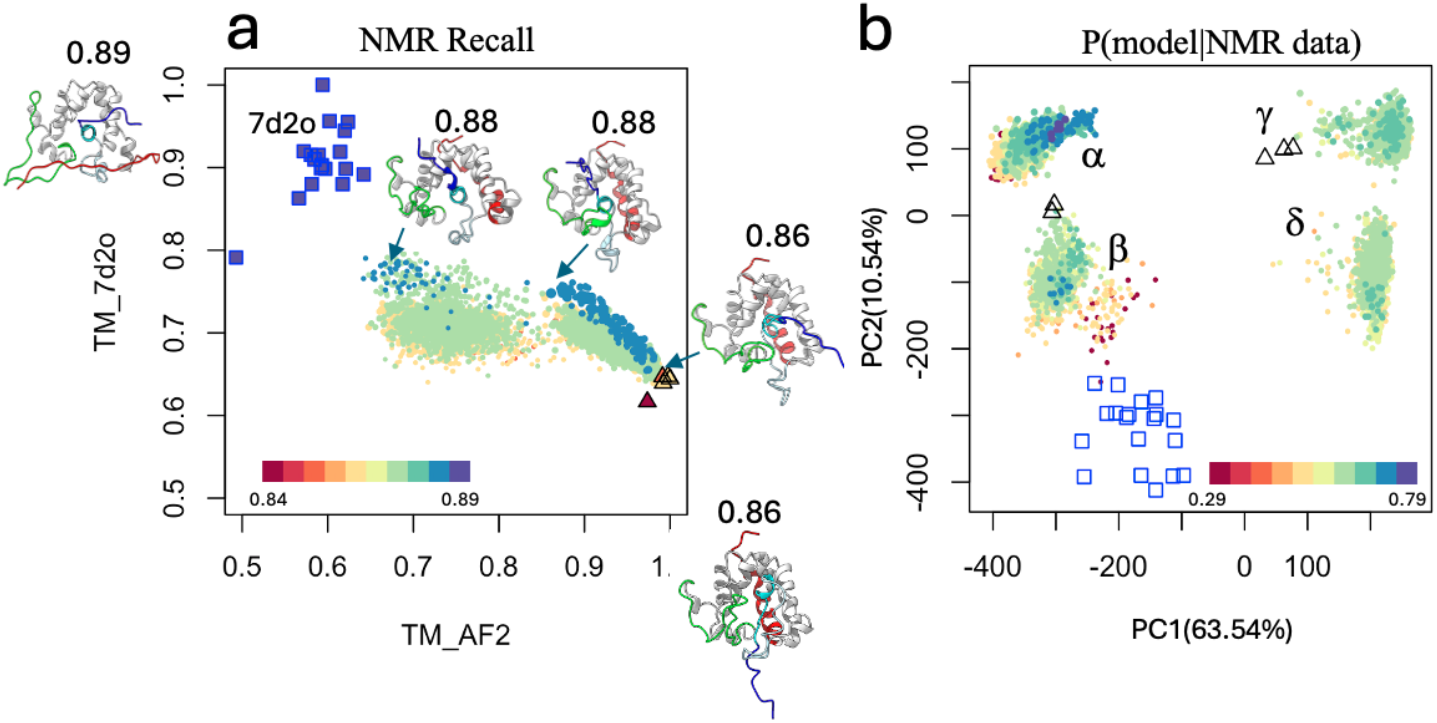
Diversity of AFsample models. (**a**) TM plots of AFsample vs NMR_7d2o_ and AF2 models. TM_AF2 scores are calculated by comparing each conformer with the standard AF2 rank1 model (for ordered regions 42-78 and 93-162). TM_NMR_7d2o_ scores are calculated by comparing each conformer with the medoid NMR_7d2o_ model (model #3, for ordered regions 7-19, 34-81, and 95-143). The dot plot for each conformer is colored by its NOESY recall score for AFsample (circles), conventional NMR_7d2o_ (squares), and standard AF2 (triangles) models. (**b**) 2D projections of C*α* coordinate principal component analysis (PCA). Models are colored by Bayesian posterior score P(model|NMR data), defined in Methods. Conventional NMR_7d2o_ models are indicated as blue open squares and standard AF2 models are indicated as black open triangles. In all panels, the size of each AFsample point is scaled by the pTM score of the model, the bigger the point the higher the pTM. The four PCA clusters (*α,β,γ*, and *δ*) are labeled in panel **b**.

Principal component analysis (PCA) grouped AFsample models into four clusters (α, β, γ, and δ; **Figure 2b**). These models were scored against NOESY and RCI data using Bayesian posterior P(model | NMR data) metrics (Methods**; Supplementary Fig. S3**). Clusters α (∼20%) and β (∼20%), with high NOESY recall scores, represented plausible conformations, while clusters γ (∼25%) and δ (∼35%) with lower recall scores were less likely. Structural differences among clusters were primarily driven by the orientation of helix H1 (PC1) and the position of the H5/H6 loop (PC2; **Supplementary Figure S4**).

### Conformer selection and identification of states

Conformational states identified by PCA were scored using experimental NMR data to calculate Bayesian posterior probability P(model ∣ NMR data) (**Eqn. 11)** metrics, enabling the selection of models from each cluster that best fit the data (Methods). Four candidate single-state ensembles (*α*5, *β*5, *γ*5, and *δ*5) were created from the top five highest-scoring models of each cluster, ranked by their average P(model | NMR data) scores. The ensemble *α*5, which had the highest P(model ∣ NMR data), was identified as state 1.

Residue-by-residue comparisons of ensemble-based backbone C*α* RMSF values (RMSF_ENS_) calculated from state 1 with experimental RMSF estimates from chemical shift-based RCI data (RMSF_RCI_) revealed discrepancies, interpreted as limitations of single-state modeling (blue line in **Figure 3a**). To improve this agreement, state 1 was combined with each of the other candidate single-state ensembles β5, γ5, and *δ*5, to create composite RMSF_ENS_ predictions, assuming fast (or fast-intermediate) exchange. Agreement with the experimental RMSF_RCI_ profile was quantified using Concordance Correlation Coefficients (CCCs) (**Eqn. 12**). For GLuc, the combination of state 1 and the ensemble β5 maximized the CCC (0.55), outperforming the other two-state combinations (0.12 and 0.06; **Supplementary Figure S5**). This combined CCC is also significantly higher than that of the original NMR_7d2o_ structure (0.30), or the standard AF2 model (0.05). Accordingly, clusters γ and δ, with flipped helix H1 orientations, were excluded due to low composite CCC values. This flipped H1 orientation was also not supported by NOESY data. The ensemble β5 was therefore identified as state 2.

**Fig. 3.**
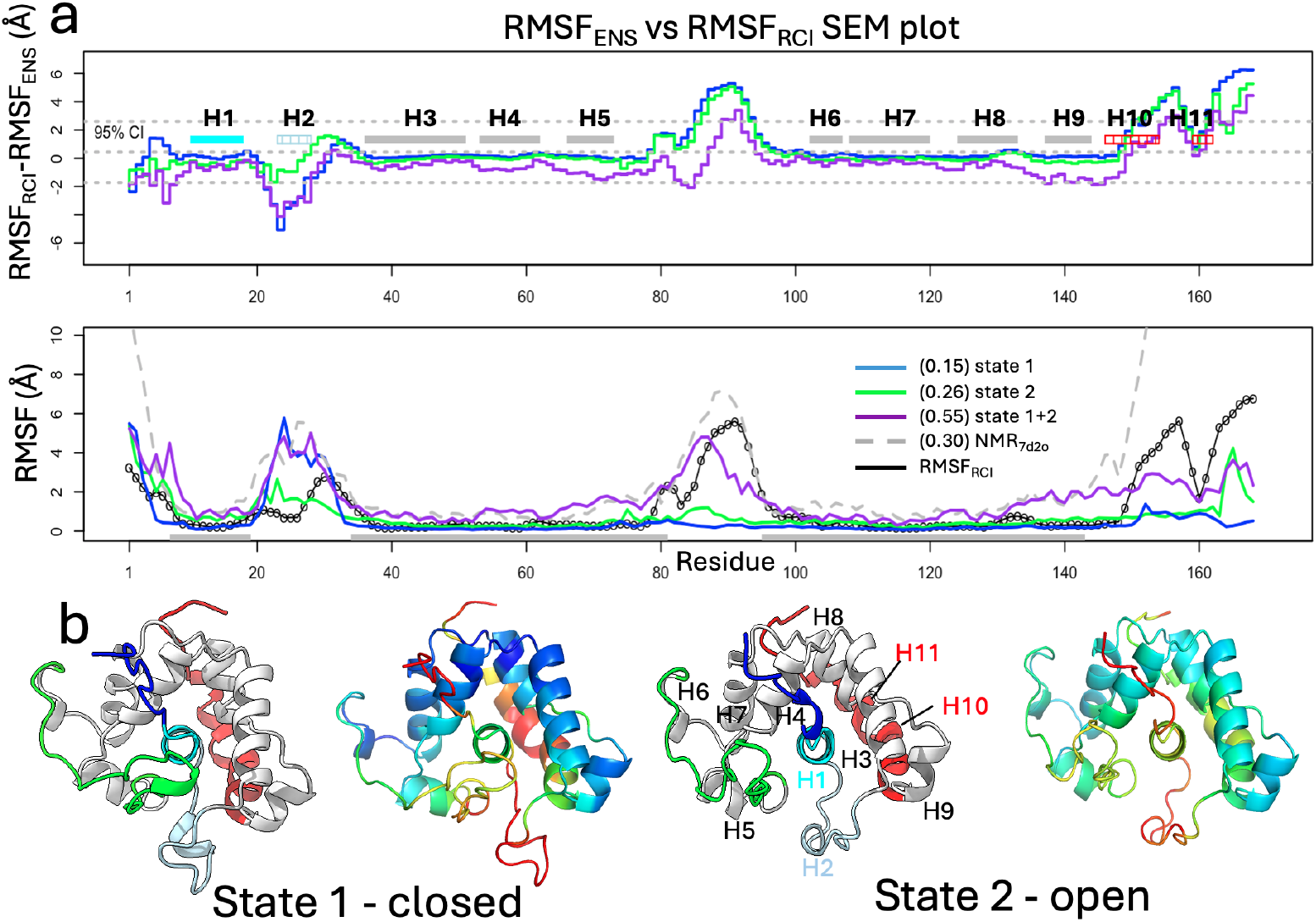
Combining states by matching RCI data with modeled atomic RMSFs. **(a)** (top) Comparison SEM plots of RMSF_RCI_ - RMSF_ENS_ for state 1, state 2, and their 1:1 mixture. (bottom) RMSF_ENS_ for state 1, state 2, their mixture, and for NMR_7d2o_ compared with RMSF_RCI_, with “well defined” NMR_7d2o_ regions shown as gray bars at the bottom and used in structural alignments for RMSF_ENS_ calculations. CCC scores for RMSF_ENS_ of selected conformers vs RMSF_RCI_ are provided in parenthesis in panel (a) inset. (**b**) Conformer-selected AlphaFold-NMR multi-state structures of GLuc. (left pair) State 1 (rank 3 model) is colored by structural elements and by per-residue pLDDT scores, respectively. (right pair) State 2 (rank1 model) colored the same.

Comparison plots of RMSF_ENS_ vs RMSF_RCI_ (**Figure 3a**), show improved agreement for the mixture of states 1 and 2 compared to either of the two single states, particularly in regions of the H5/H6 loop, and the C-terminal helix H10/H11. Representative structures of the “closed” (state 1) and “open” (state 2) conformations are shown in **Figure 3b**. In particular, the RMSF_ENS_ fluctuations for the H5/H6 loop (“lid”) align more closely with the RMSF_RCI_ data for the combined state 1 and 2 models than for individual states.

The AF-NMR ensemble, consisting of ten models from combined states 1 and 2, was further assessed by comparing per-residue pLDDT values with NMR RCI data (**Supplementary Figure S6**). Each state exhibited distinct pLDDT profiles, with per-residue average pLDDT (pLDDT_avg_) values for states 1 and 2 showing improved correlation with experimental RCI data compared to standard single-state AF2 models (improved SCCs, **Supplementary Figure S6**), especially for helix H1. Most residues agree within the 95% confidence interval except the H5/H6 loop, which requires a combined model for accurate representation. Using pLDDT_avg_ values as proxies for *state-specific conformational flexibility*, helices H3– H5 and H6–H9 have low flexibility in both states. Helix H2 and the H5/H6 loop are more flexible in both states, while helix H1 exhibits greater flexibility in state 2, and the C-terminal helices H10 and H11 are more flexible in state 1.

### Cross-validation of conformational states by NOESY Double Recall analysis

NOESY Double Recall analysis confirmed that the combination of states 1 and 2 better explains the NMR NOESY data than single-state models, including NMR_7d2o_ (**Figure 4a; Supplementary Figure S7 and Table S1**). State 1 uniquely explains 185 NOEs (101 long-range) compared to NMR_7d2o_, while state 2 uniquely explains 164 NOEs (71 long-range). The combined ensemble of states 1 and 2 accounts for 218 NOESY peaks (116 long-range) that are not consistent with the NMR_7d2o_ structure, confirming that this two-state ensemble fits the NOESY data more effectively than the NMR_7d2o_ model. However, a small number of NOESY peaks are not explained even by the combined ensemble, suggesting additional states may exist in dynamic equilibrium (**Supplementary Table S1**). Additionally, Double Recall analysis was used to identify distinct subsets of NOEs for each state (**Figure 4a-d**) that demonstrate the ability of AF-NMR to resolve multiple conformational states and capture dynamic transitions essential for GLuc’s function.

**Fig. 4.**
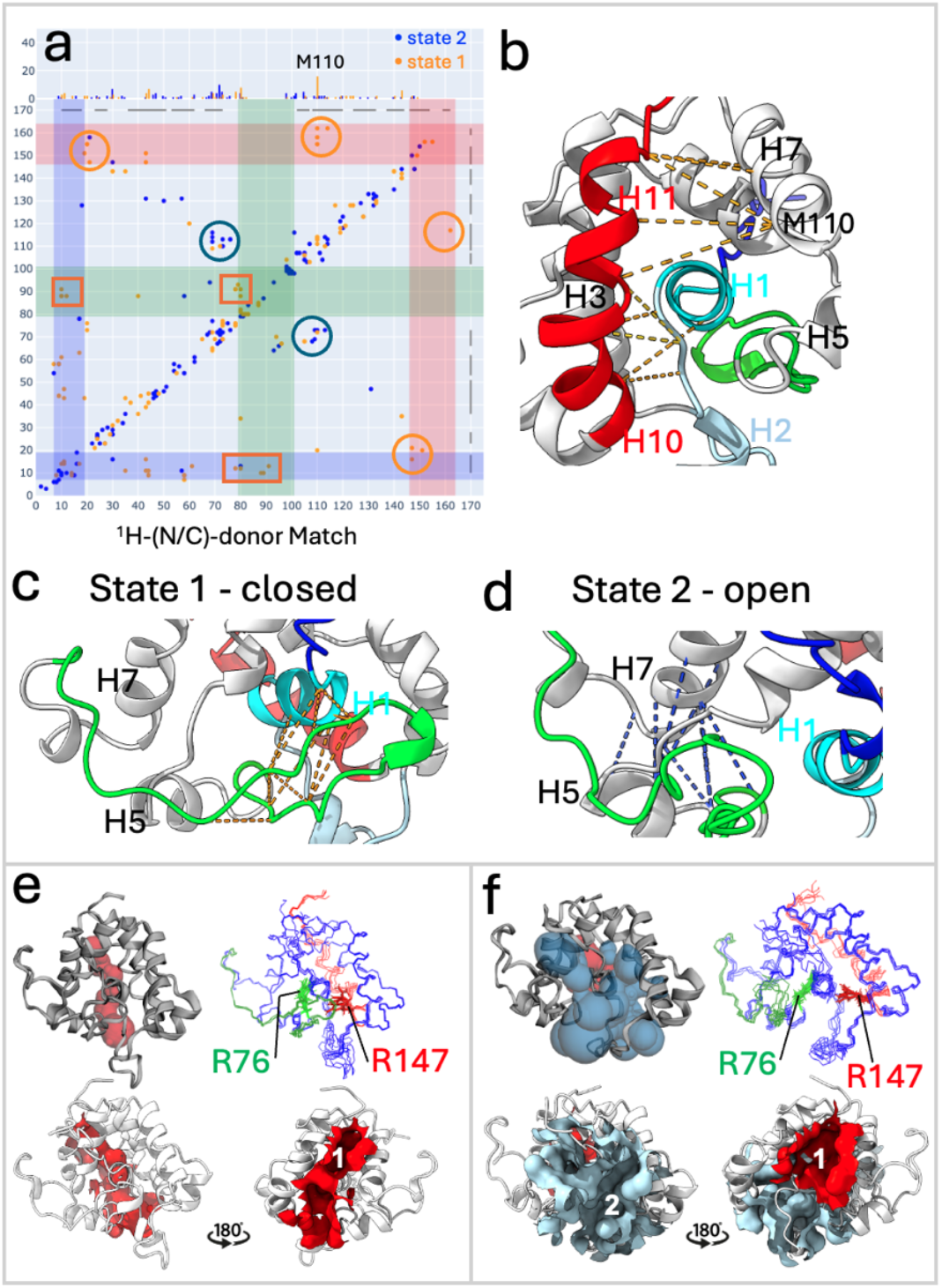
Detailed analysis of conformer-selected AlphaFold-NMR multi-state structures of GLuc. (**a**) Double Recall plot of AF-NMR state 1 vs state 2, illustrating key inter-residue contacts distinguishing these states; orange dots support “closed” state 1, blue dots support “open” state 2, and brown dots are overlapped blue and orange dots. Scaling of the top inset is the same as for **Figure 1b**. (**b**) “Closed” state 1 with key NOEs indicated as orange dashed lines (corresponding to orange dots in the red shaded region of panel **a**) includes transient helices H10 and H11 (red) which contribute to forming part of pocket #1. (**c**) “Closed” state 1 with key NOEs indicated as orange dashed lines (corresponding to orange dots in the green shaded region of panel **a**). (**d**) “Open” state 2 with key NOEs indicated as blue dashed lines (corresponding to blue dots in the blue circles of panel **a**). In panels **b, c**, and **d**, the dashed lines connect the C*α*-C*α* atoms of the corresponding residues. **(e)** “Closed” state 1 includes a pocket (pocket #1, red), while **(f)** “open” state 2 has both a larger pocket #1 and a new pocket (pocket #2, blue). In the “closed” state 1, the H5/H6 loop contacts helix H1 closing binding pocket #2, while in the “open” state 2, this loop moves close to helices H5 and H7, opening binding pocket #2. The pocket of “closed” state 1 (red pocket #1, shown for state 1, rank 3 model with highest pTM score in cluster *α*) has a pocket volume of 106 Å^3^, while “open” state 2 includes not only a wider red pocket #1 (shown for state 2, rank 1 model with the highest pTM score in cluster *β*, with volume 173Å^3^), but also a new large cryptic blue pocket #2, volume 786 Å^3^. Side chains of functionally-important residues Arg76 (green) in pocket #1 and Arg147 (red) in pocket #2 are shown in state 1 and state 2, respectively (panels **e**,**f** – upper right).

### Conformational ensembles and structural quality assessment

The conformational ensembles for states 1 and 2 have been deposited in the Protein Data Bank (PDB ID: 9A8V). These ensembles exhibit significantly better structure quality metrics compared to conventional NMR structures 7d2o and 9fla (**Supplementary Tables S2–S5**). NOESY RPF scores, which compare global NOESY peak list data with structural models, are similar for state 1, state 2, and NMR_7d2o_ (**Supplementary Table S6**); NOESY data for 9fla are unavailable.

### Structural features of GLuc states 1 and 2

States 1 and 2 represent two substantially-populated alternative conformations of GLuc in fast-exchange (or fast-intermediate-exchange) dynamic equilibrium. A detailed structure-function analysis (**Figure 4b-f**) reveals that the major difference between these states lies in the conformation of the H5/H6 thumb-shaped loop. In the “closed” state 1, the loop forms a “lid” capping a surface binding pocket (**Figure 4c**,**e**), supported by ∼8 NOEs (residues 88–91 and 10–16; **Figure 4a**). In the “open” state 2, the loop moves away, exposing a cryptic pocket and allowing helix H5 to interact with helix H7 (∼15 NOEs; **Figure 4d,f**). Small changes in interhelical angles (H3–H4 and H7–H8) accompany this transition, with state 1 forming a compact “pincher” structure and state 2, having an expanded pocket. Helix H2 transitions from an *α*-helix in state 1 to a 3_10_ helix in state 2. Both state 1 and state 2 include a C-terminal “broken helix”, defined by two helices H10/H11. This broken helix is packed closer to the helices H1 and H7 in state 1 than in state 2. Specifically, in state 1 helices H10/11 have ∼12 NOEs interacting with helix H7 (including 5 NOEs to Met110) and the end of helix H1 (**Figure 4a,b**). Residue Met110 exhibits several NOEs unique to state 1 (**Figure 4a**) indicating its potential role in modulating the two-state equilibrium.

The transition between states also reshapes two surface pockets of GLuc (**Figure 4e,f**). In state 1, pocket #1 is small and variable (∼30 – 110 Å^3^), while in state 2 pocket #1 expands (∼ 170Å^3^) and a new larger cryptic pocket #2 appears (∼600–800 Å^3^). Residues Arg76 (in pocket #1) and Arg147 (in pocket #2) both have critical roles in GLuc bioluminescence^32,51^, and these structural changes likely impact substrate binding and enzymatic function (see Discussion).

### Conformational switching of CDK2AP1

As a second example, the AF-NMR method was also applied to investigate the dynamic structure of CDK2AP1, a homodimeric right-handed four-helical bundle^47^.

Conventional NMR and standard AF2 models of CDK2AP1(residues 61–115) are similar (RMSD: 0.99–1.21 Å) but neither position Cys105 sulfur atoms within 3 Å for disulfide bond formation, which has been suggested (but not demonstrated) to be important for CDK2 binding^48^. Using AF-NMR conformational selection (**Supplementary Figure S2)**, 6000 AFsample models were analyzed and grouped into four clusters (α, β, γ, and δ; **Figure 5a)**. Most models belong to clusters *α* (∼50%) and *β* (∼47%), while clusters *γ* (∼2%) and δ (∼1%) represent alternative packing arrangements (**Supplementary Figure S8**) that are excluded under the conditions of the NMR study by NOESY data.

**Fig. 5.**
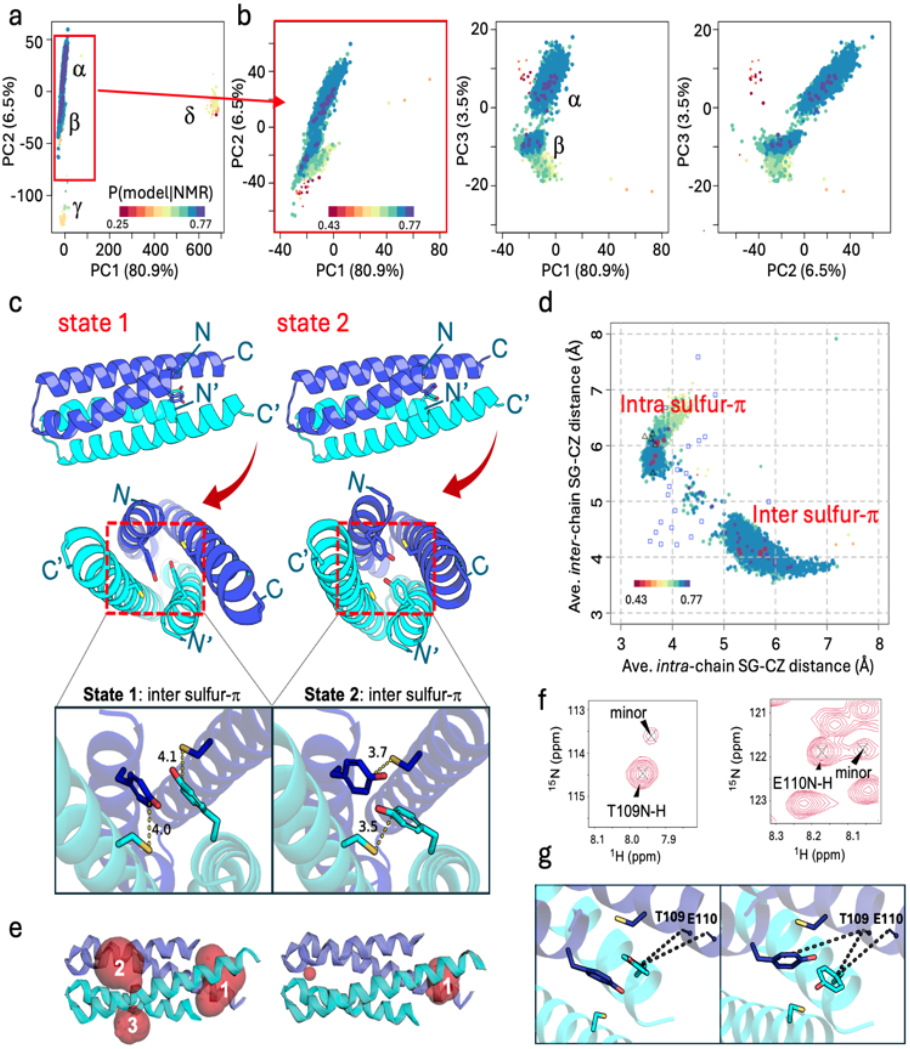
Hidden conformational states of CDK2AP1. **(a)** PCA based on the C*α*-C*α* distance matrix, showing structural variation along PC1 and PC2. Clusters (*α, β, γ*, and *δ*) correspond to distinct conformational states. **(b)** Expanded PCA plots for clusters *α* and *β* (red rectangle in (a)). Clusters *α* and *β* are well separated by PC3. Model PCA coordinates are colored based on values of P(model | NMR data), defined in the inset of panels a,b. The size of each point is coded by the pTM score of the model, the larger the point the higher the pTM. **(c)** Ribbon diagrams illustrating inter S-*π*state (state 1, rank 2) with key S*γ*-Cζinter chain distances (4.0 and 4.1 Å^2^) and intra sulfur-πstate (state 2, rank 1) with S*γ*-C*ζ*intra-chain distances (3.5 and 3.7 Å^2^). Side chains of Tyr63 and Cys105 are displayed in stick representation with corresponding distances labeled. **(d)** Scatter plot of average intra-chain and inter-chain distances of Tyr63-C*ζ* to Cys105-S*γ* for all models from clusters α and β. The top 5 models from clusters α and *β* are circled in red. Conventional NMR models (PDB_ID 2kw6) are indicated as blue open squares and standard AF2 models are indicated as black open triangles. **(e)** Surface pockets identified by CASTpFold^52^ (**Supplementary Figure S14)**. The state 1 model with the largest pockets (#1-#3) is shown. The state 2 model has a smaller pocket #1, while pockets #2 and #3 were each < 2 Å^3^. These surface cavities are shown in translucent red. **(f)** ^15^N SOFAST HMQC spectral regions showing peaks of T109 and E110 HN corresponding to major and minor states. **(g)** NHs of T109 and E110 are shown as sticks with close distances (< 7.5 Å) to Tyr63 rings indicated with dashed lines.

The AFsample models were scored against experimental NMR data to estimate P(model | NMR data) (**Figure 5a,b**; **Supplementary Figure S9**). The five top-scoring conformers from clusters *α* and *β* were identified as states 1 and 2, respectively. Comparison of RMSF_ENS_ vs RMSF_RCI_ data shows improved agreement when states 1 and 2 are combined (CCC = 0.70), outperforming single-state models (CCC = 0.39 for state 1; CCC = 0.14 for state 2; **Supplementary Figure S10**). This plot also reveals greater flexibility in the C-terminal region of helices H2/H2’ in state 1 compared to state 2, which is also reflected in the lower state-specific pLDDT scores (**Supplementary Figure S11**). The two-state model for CDK2AP1 was further validated by NOESY Double-Recall analysis, which identified peaks specific to each state (**Supplementary Figure S12 and Table S7**). However, as the structures of these two states are very similar (**Figure 5c,d**; pairwise backbone RMSDs: 0.56 - 1.32 Å) and the spectra of these homodimeric conformers are highly degenerate, only a few NOESY peaks are unique to one or the other conformational states. Of the observed NOEs, 22 (16 long-range) were unique to state 1, while 23 (3 long-range) were unique to state 2. While most chemical shifts do not distinguish the two states, peak doubling of Thr109 and Glu110 HNs, which experience ring-current shifts, indicates slow exchange between these major (∼85%) and minor (∼15%) states (**Figure 5e,f,g**; **Supplementary Figure S13**).

None of the cluster *α* and *β* conformers have Cys105 sulfur atoms with distances suitable for disulfide-bond formation (< 3 Å); however, favorable geometries for sulfur – aromatic π interactions between Cys105 and Tyr63 were observed. These interactions, along with differences in surface pockets, distinguish states 1 and 2 (**Figure 5c-e**). In state 1, *inter*-chain sulfur – π interactions and Tyr63 π-π stacking lead to a more open structure between the N- and C-terminal ends of the four helices, forming an ‘end’ cavity (pocket #1). Several residues in this pocket exhibit peak doubling due to slow conformational exchange (**Figure 5e, Supplementary Figures S13 and S14**). Pockets #2 and #3 contain multiple basic residues (Lys and Arg). In contrast, state 2 features *intra*-chain sulfur - π interactions, leading to a more compact structure with a smaller pocket #1, while pockets #2 and #3 are significantly reduced or absent (**Figure 5c and e, right, Supplementary Figure S14)**. Upfield shifts in the Cys105 H*β* protons support both intra- and inter-chain sulfur–π interactions due to ring-current effects (Ringer, 2007). The conventional NMR structure (2kw6) captures elements of both states, but does not fully represent either due to conformational pinning^53^; its intra- vs inter-chain Cys105 S*γ* - Tyr63 C**ζ** distances fall between the two states (**Figure 5d**). Standard AF2 consistently generates only state 2, favoring the more compact structure.

## Discussion

### Conformational selection with AF-NMR

The AF-NMR method combines AlphaFold modeling with enhanced sampling and NMR-guided conformational selection, offering a novel approach to protein structure determination. By generating diverse conformers and scoring them against NMR data, including NOESY recall and RCI metrics, AF-NMR identifies single or multiple conformational states that best fit the experimental data. Alternative states are further validated using NOESY Double Recall analysis. Unlike conventional restraint-based NMR modeling, which relies on distance constraints to build a single-state model, AF-NMR applies a Bayesian framework to integrate sample-specific NMR data with AF2 pTM and per-residue model reliability scores (pLDDTs) allowing for selection of one or more conformational states that together best fit the NMR data.

Conventional restraint-based NMR approaches can result in inaccurate models, particularly when dealing with conformationally heterogeneous systems. Sparse NMR data from transiently populated states may be insufficient for accurate modeling, while conflicting NOEs arising from multiple states can result in partially satisfied restraints or overly constrained structures. These issues can lead to conformational “pinning,” where the requirement to simultaneously satisfy distance restraints arising from distinct states forces the model into an unrealistic compromise^53,54^. Difficulties also arise from peak degeneracy and assignment of ambiguous NOEs in flexible regions. In contrast, AF-NMR allows for the explicit selection and combination of multiple conformational states without distortions from restraints, leveraging AI-based modeling to generate candidate structures that align with the experimental data.

Consistent with other studies^1,9-13,49^, we observe that AF2 residue-specific pLDDT scores reliably distinguish ordered from disordered regions. However, discrepancies arise in regions where pLDDT does not correlate with experimental flexibility metrics, such as RCI and hetNOE. Here we compare RCI with pLDDT of standard AF2 models, an estimate of the local rigidity (or certainty). We reasoned that discordance between these metrics can be an indication of conformational diversity not captured by the standard AF2 models. Using enhanced sampling protocols, such as AFsample, AF-NMR generates more diverse conformers and improves the nonlinear agreement between pLDDT and RCI (i.e., |SCC(pLDDT, RCI)|). In cases where single-state models fail to fully explain the experimental data, combining multiple state models enhances agreement, providing a more accurate representation of dynamic structural ensembles.

Conformational state bias in AlphaFold modeling has been increasingly recognized, with several studies showing how memorization of states in training data can bias predictions of proteins with alternative native structures^35,37,38^. If experimental structures which capture only one (or some) of the states were included in its training set, conformational state bias may prevent AF from finding alternative states.

While AF2 was not trained on NMR structures^1^, state bias may still occur if X-ray or cryoEM homolog structures of the target protein were included in its training set. In this study, we avoided such bias by selecting proteins with no homologous structures in the training set. However, where bias is suspected, alternative sampling strategies, such as varying MSA inputs or even retraining AlphaFold with specific exclusions may be necessary to ensure unbiased sampling.

Although not yet broadly adopted by the structural biology community, we and others have emphasized the potential value of Bayesian formalisms in NMR data analysis, particularly for integrative modeling (Andrec *et al*., 1999; Andrec *et al*., 2000; Hoch and Stern, 2005; Rieping *et al*., 2005; Nilges *et al*., 2008; Rieping *et al*., 2008). The Bayesian framework used in AF-NMR can support incorporation of diverse NMR data types, including residual dipolar coupling, back-calculated chemical shift, chemical exchange, nuclear relaxation, paramagnetic relaxation enhancements, and pseudocontact shift data, offering a comprehensive toolset for conformer selection and validation. As an evolving method, AF-NMR may also benefit from integrating alternative AI-based modeling methods such as AlphaFold3^55^, RosettaFold^56^, or others^57,58^. Incorporating alternative AI-based enhanced sampling techniques^17^ will also further expand AF-NMR’s ability to capture conformational heterogeneity and dynamics.

### Dynamic Structural Features of GLuc

AF-NMR reveals distinct conformational states of GLuc that are consistent with previous structure-function studies^28,31,32,51^ but provide additional insights into its structural transitions. The H5/H6 loop (“lid”) undergoes a dynamic shift between a “closed” state 1, where it caps pocket #2, and an “open” state 2, where the cryptic pocket #2 is exposed. This transition reshapes GLuc’s surface cavities, which house residues critical for bioluminescence, including Phe72, Ile73 and Arg76 in pocket #1 and Trp142, Leu144, Arg147, and Phe151 in pocket #2. The interplay between these pockets in its structural transition between states is consistent with GLuc’s reported positive cooperativity in coelenterazine substrate binding, characterized by a Hill coefficient of ∼1.8^59,60^. Substrate binding at one pocket may enhance affinity at the other, facilitated by the structural changes observed during the closed-to-open transition. Additionally, the C-terminal helices H10 and H11 exhibit dynamic behavior, alternating between an ordered “broken helix” (in state 2) and a less ordered conformation (state 1), as indicated by pLDDT profiles (**Supplementary Figure S6**). These dynamic transitions are consistent with intermediate exchange broadening observed in ^15^N relaxation data^28,31^. Together, these findings highlight how conformational changes in GLuc shape its functional binding pockets and regulate its bioluminescence activity.

### Dynamic Structural Features of CDK2AP1

AF-NMR reveals two distinct conformational states of CDK2AP1 in dynamic equilibrium. In state 1 inter-chain sulfur– π interactions between Cys105 and Tyr63 contribute to the interface between helices H2 and H2’. In contrast, in state 2 these interactions switch to intra-chain sulfur–π interactions, resulting in a more compact structure. Sulfur–π interactions, although weaker than traditional hydrogen bonds or π–π stacking, still significantly contribute to protein stability and are frequently observed in protein structures, predominantly with sulfur positioned above the aromatic ring face (Ringer, 2007) as observed here for CDK2AP1.

The dynamic equilibrium between states 1 and 2 is supported by RMSF_ENS_ vs RMSF_RCI_ comparisons and NOESY Double Recall analysis, which identified unique NOEs for each state. Doubled resonances for Thr109 and Glu110 HNs further validate multiple states, reflecting distinct aromatic ring current shifts. Slow exchange between these states indicates that the two conformations of CDKAP1 interconvert on a timescale longer than milliseconds and that NOEs for the many other degenerate resonances of these two structurally-similar conformational states are overlapped (**Supplementary Figure S13**).

Although the conventional NMR structure (PDB: 2kw6) is high quality and fits the NOESY data well^47^ (**Supplementary Table S11**), it does not fully capture the distinct states identified by AF-NMR. Some NMR models have short *intra*-chain S*γ*–Cζ distances (<4 Å), but none exhibit short *inter*-chain S*γ*–Cζ distances (< 4 Å) or full Tyr63 π–π stacking. Rather, these structures are distorted by conformational pinning^53^, an artifact of interpreting NOEs arising from multiple states to restrain a single model. As a result, *inter*-chain S–π interactions were absent, while *intra*-chain S–π interactions (state 2-like) were overlooked. Additionally, the dynamic transition of cryptic pocket #3 from a closed state (state 2) to an open state (state 1), along with the identification of pockets #1 and #2 (**Supplementary Figure S14**), was not resolved in the restraint-based NMR models. These findings underscore the advantage of AF-NMR in capturing conformational heterogeneity and cryptic pockets.

The dynamic remodeling of CDK2AP1’s pockets in the transition from state 2 (closed) to state 1 (open) may be functionally relevant for CDK2 binding and other interactions. Although Cys105 is critical for CDK2 interactions^48^, mutation of Cys105 to Ala does not significantly disrupt CDK2AP1’s dimeric structure ^47^. Ablation of Cys105 - Tyr63 interactions is, however, expected to affect the dynamic equilibrium between states 1 and 2. Additionally, residues Thr109, Glu110, and Arg111 in pocket #1, which expands in the open state, has been proposed as a key CDK2 binding site in HEK-293 cells^39^. Beyond CDK2 binding, the positively charged residues in pockets #2 and #3 may facilitate interactions with other reported binding partners, such as negatively charged regions at the C-terminus of CHD3 and CHD4 in the NuRD complex^44,45^. Since pocket #3 is only present in state 1 (**Supplementary Figure S14)**, a transition from a closed to an open state may regulate accessibility to these interaction sites. These findings suggest that state-dependent pocket formation could modulate CDK2AP1’s binding properties, influencing both CDK2 regulation and interactions with other partners.

## Conclusions

In this study, we applied AFsample^21,50,61^, an enhanced conformational sampling method originally developed for improving model predictions of protein complexes to generate diverse conformational states of GLuc and CDK2AP1. Using a Bayesian selection metric, models were selected based on their agreement with global NMR NOESY (*recall* score) and residue-specific chemical shift RCI data. Where AF-NMR revealed alternative conformational states, these were rigorously cross-validated using atom-pair-specific NOESY data with Double Recall analysis. Our findings underscore the utility of modern AI-driven deep learning methods in generating realistic conformational states, including accurate models that can be identified and validated using experimental data. *Enhanced sampling methods proved invaluable in capturing structural heterogeneity and identifying regions of flexibility and rigidity that align with experimental RCI, NOE, and nuclear relaxation data*.

Beyond the systems studied here, AF-NMR has the potential to be broadly applied to investigate conformational heterogeneity and dynamics in a wide range of proteins, including proteins and complexes undergoing both large- and small-scale structural transitions. Multiple conformational states that can be more accurately captured by AF-NMR with enhanced sampling are particularly evident in regions of proteins where standard AF2 pLDDT values are high (i.e. high single-state model confidence), yet experimental chemical shift or nuclear relaxation data reveal underlying conformational variability. This is commonly observed in NMR studies of small proteins. By combining AI-based sampling and NMR-guided conformer selection, AF-NMR offers a powerful tool for studying protein dynamics and uncovering structural heterogeneity that cannot be resolved using conventional methods. These AF-NMR models provide a detailed framework for understanding these “hidden” states and their functional implications. The incorporation of additional experimental data types, such as residual dipolar couplings (RDCs), paramagnetic relaxation enhancements (PREs), or pseudocontact shifts (PCSs) and/or integration of emerging AI-based modeling methods, such as AlphaFold 3 or OpenFold, and advanced sampling techniques will further expand AF-NMR’s applicability to more challenging systems, including large multimeric complexes and transiently populated states. These studies demonstrate how enhanced sampling combined with NMR data can uncover previously unrecognized structural and dynamic features, including cryptic pockets and alternative conformations, offering a powerful and evolving tool for studying protein dynamics, molecular recognition, and function.

## Methods

### Experimental NMR data

The published NMR structure of GLuc (PDB ID 7d2o) was determined using extensive ^1^H,^13^C and ^15^N NMR assignments, along with ∼2500 NOE-based distance restraints, including ∼580 long-range restraints^28^. Chemical shift assignments were obtained from the BioMagResDatabase (BMRB ID 36385). NOESY peak lists and HetNOE data were provided by Wu, Kobayoshi, Kuroda, Yamazaki, and colleagues, who originally determined the NMR structure used for CASP14 structure prediction assessments^28,29^, and are archived at https://zenodo.org/records/13831427.

The solution NMR structure of CDK2AP1 (PDB_ID 2kw6) was determined previously using standard triple-resonance NMR and 3D NOESY data, including 3D X-filtered NOESY data on a mixed sample of 50% [^15^N, ^13^C]-labeled and 50% unlabeled CDK2AP1 to resolve key inter-chain dimer NOEs^47^. NOESY peak lists and updated chemical shifts from BMRB ID 16808 (including newly assigned H58 - K62 resonances) were used for RPF analysis and AF-NMR model selection. To generate simulated NOESY peaks for AF-NMR states 1 and 2, we used *simulateNOE*.*pl*, a program developed for NMR-assisted CASP13 assessments^62^. The simulated peak lists were analyzed using Double Recall; peaks unique to each conformer were identified and used to guide further peak picking of weaker peaks at lower contours. An additional 124 NOESY peaks were picked by this iterative NOESY peak list refinement and used in the final Double Recall analysis [**Supplementary Table S7;** refined NOESY peak list available at (https://github.rpi.edu/RPIBioinformatics/AlphaFold-NMR)].

### AlphaFold2 modeling

AF2 models were generated using the *ColabFold* server of AF-multimer (version 1.5.3) using default setups with no templates^63^. GLuc’s ColabFold models were nearly identical to DeepMind’s CASP14 AF2 models^29^. No X-ray crystal structures of GLuc or CDK2AP1 homologs were available in the PDB at the time of training AF2. The top five-ranked relaxed AF2 conformers included hydrogen atoms and per-residue pLDDT scores, as generated by the ColabFold pipeline.

### AF2 enhanced sampling with AFsample

Enhanced sampling with AFsample was conducted as described by Wallner^21,50,61^ using a modified *AlphaFold v2*.*2*.*0* pipeline^64^ (https://github.com/bjornwallner/alphafoldv2.2.0), installed on the AiMOSx NPL cluster in the Center for Computational Innovation at Rensselaer Polytechnic Institute. The AF2 training set (PDB database from April 30, 2018) excluded NMR structures^1^. AFsample calculations, which use six different distinct dropout settings^21,50,61^ to generate ∼6000 diverse structural models. Hydrogen atoms were added using *run_relax_from_results_pkl*.*py* from the AFsample package. AFsample is quite aggressive in generating conformational diversity, and computes some models that are not physically reasonable, with incorrect amino acid chirality, non-native cis peptide bonds, missing residues, and other biophysically incorrect features, particularly in the not-well-packed residue segments. The most egregious of these physically unreasonable models were identified and removed, as described elsewhere (Spaman et al, manuscript in preparation). Final datasets included 4,990 relaxed models for GLuc and 5984 relaxed models for CDK2AP1. The pTM scores were retrieved from AF2 output files (*result_model_\**.*pkl*.*json*), and pLDDT scores from atomic coordinate files.

### Knowledge-based protein structure validation

All model quality assessments were performed using the *Protein Structure Validation Software* (*PSVS* v2.0)^65^ (https://montelionelab.chem.rpi.edu/PSVS/PSVS2/) and the protein model assessment program *PDBStat* (Tejero *et al*., 2013). *PSVS* runs a suite of knowledge-based software tools including *ProCheck* (v3.5.4) (Laskowski *et al*., 1993) and *MolProbity* (v6.35.040409)^66^. Validation scores were normalized as Z-scores, referenced against mean values obtained for 252 high-resolution (<1.8 Å) X-ray structures. For all metrics assessed, positive Z scores indicate ‘better’ scores^65^.

### NMR NOESY RPF-DP scores

RPF-DP scores provide a rapid and sensitive assessment of how well a structural model agrees with 3D ^15^N and ^13^C-resolved NOESY peak lists and resonance assignments, serving as a measure of structure accuracy^29,67,68^. These scores were computed using the *RPF server* (ASDP ver 1.0) https://montelionelab.chem.rpi.edu/rpf/. *RPF* scoring includes several key metrics: *recall*, the fraction of NOESY peaks explained by the model (or collection of models); *precision*, the fraction of short interproton distances in the model(s) supported by NOESY peaks; *F-measure*, the geometric mean of recall and precision; and *discriminating power* (*DP)*, a normalized F-measure that accounts for NOESY data completeness and the expected baseline for a random coil structure^29,67,68^. RPF-DP scores serve as an NMR equivalent to an “R-factor”, allowing model evaluation against unassigned NOESY peak list data. Output folders (zip files) were downloaded from the *RPF server* for further Double Recall analysis (described below).

### NMR NOESY Double Recall analysis

Double Recall analysis (v1.0) was performed in a Jupyter notebook to compare two conformer ensembles (A vs B) and identify NOEs uniquely explained by each. The method uses *RPF* recall violation reports from zipped RPF output files to determine NOEs that correspond to short inter-proton distances present in only one ensemble. NOEs unique to ensemble A are plotted in blue, while those unique to ensemble B are in orange. The results are displayed in a contact map-like 2D plot, where the x-axis represents the residue number of the first H(-N/C) proton (“1H(-N/C)-donor Match”) and the y-axis represents the residue number of the second proton (“1H-acceptor Match”). Multiple NOEs between the same residue pairs appear as a single dot. A 1D histogram at the top of the plot shows the per-residue count of uniquely matched NOEs along the x-axis, which have higher-confidence assignments due to the additional frequency resolution provided by the attached ^15^N or ^13^C nucleus.

Double Recall analysis allows for comparison of any two conformer ensembles by evaluating their midrange interproton distances for the NOESY peak comparison. The method was further extended to compare combined multi-state ensembles against a reference ensemble by using the shortest midrange interproton distances across merged conformational states. In either case, the number of models in each conformer ensemble does not need to be identical for a valid comparison.

### Well-defined residue ranges and TM scores to reference structures

Well-defined residue ranges for NMR and AF2 ensembles were determined using Cyrange^69^ and applied to structural superimpositions, quality assessments, and TM score calculations^70^. For TM-score comparisons, the medoid conformer^71^ of the NMR_7d2o_ ensemble was used, with the corresponding well-defined residues. Similarly, AF2 TM-scores were calculated using the rank1 conformer, restricted to its well-defined residues.

### Per-residue Random Coil Index (RCI) and RMSF_RCI_

RCI values were computed using the Wishart server^72^ (http://www.randomcoilindex.ca/cgi-bin/rci_cgi_current.py), applying default options:

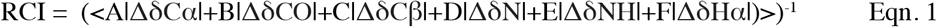

where |ΔδCα|, |ΔδCO|, |ΔδCβ|, |ΔδN|, |ΔδNH| and |ΔδHα| are the absolute values of the conformation-dependent chemical shifts ΔδX = δX_observed_ - δX_random_coil_ (in ppm) of Cα, CO, Cβ, N, NH and Hα resonance, respectively, and *A, B, C, D, E* and *F* are weighting coefficients. Left angle and right angle brackets (<·>) indicate that the RCI per residue is the average value across the several resonances (^1^H, ^13^C, and ^15^N) of that residue. If any per-residue RCI value exceeds 0.6, it is replaced with the ceiling limit of 0.6. After this, a second smoothing along the sequence of RCI values is done using three-point averaging to obtain the final RCI values as previously described^72^.

Root-mean-square-fluctuations based on RCIs (RMSF_RCI_), were calculated as previously described^72^:

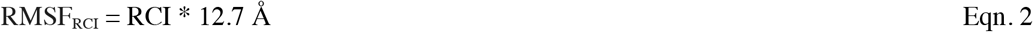

### Comparison SEM plots: pLDDT vs RCI and RMSF_ENS_ vs RMSF_RCI_

RCI values were scaled to the 0.0-1.0 range (RCI_0.6_ = RCI/0.6) for comparison with pLDDT scores. For pLDDT vs RCI plots, per-model per-residue pLDDT, ensemble-average per-residue pLDDT (pLDDT_avg_) and RCI_0.6_ values were plotted along the sequence to assess correlation. In most cases we have examined, RCI and pLDDT (or pLDDT_avg_) have an inverse correlation. The comparison plots were constructed by summing RCI_0.6_ and pLDDT_avg_ values per residue, with global mean and standard deviation (SD) calculated. Confidence intervals (95% CI) were determined using the CI t-test with one degree of freedom:

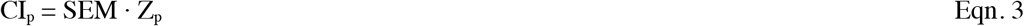

where the standard error of the mean, SEM = SD / (N -1)^½^. N is the number of residues with pLDDT > 50 and available RCI values, and Z_p_=12.71 (p=95%). Mean and CI values are displayed as dashed lines highlighting significant deviations. Residues with pLDDT ≤ 50 were excluded from the mean, SD, and CI calculations.

Similarly, for RMSF_ENS_ vs RMSF_RCI_ comparison SEM plots, atomic coordinate ensemble RMSF (RMSF_ENS_) values and RMSF_RCI_ (Eqn. 2) values were plotted along the sequence, with differences between RMSF_RCI_ and RMSF_ENS_ reported per residue. RMSF_ENS_ was computed using Bio3D^74^ in R.

### Statistical Correlation Coefficients

Spearman Correlation Coefficient (SCC) and Lin’s Concordance Correlation Coefficient (CCC) were calculated with R as described elsewhere^73^. The CCC is the concordance between a new test or measurement (e.g. RMSF_ENS_) and a standard test or measurement (e.g. RMSF_RCI_). It obeys the inequality −1 ≤ −|PCC| ≤ CCC ≤ +|PCC| ≤ +1, where PCC is Pearson’s correlation coefficient.

### Principal Component, Clustering, and RMSF analysis

To examine structural diversity across various models, principal component analysis (PCA) was conducted using Bio3D^74^. PCA was applied to C*α*-C*α* distance matrices to capture the primary structural variations among the models. Agglomerative hierarchical clustering^75^ was employed to group similar conformers into well-separated distinct clusters (states) based on PC1, PC2, and PC3.

### Bayesian model selection metric using NOE and RCI data

To quantify agreement between computational models and experimental data, an AF-NMR Bayesian model selection metric was developed, incorporating both NOE recall and RCI correlation scores. This approach estimates the likelihood of a model given NOE and RCI data using Bayes’ theorem.

To estimate how well the model fits the NOE data, the Bayesian **P(model** | **NOE) score** uses the scaled NOESY recall scores:

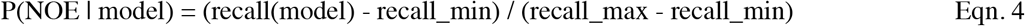

where recall_max and recall_min are the highest and lowest NOESY recall scores across all models.

The overall model probability, P(model), is estimated using the AF2 pTM metric^1^, a measure of overall structural confidence/reliability. We decided to use the pTM, rather than <pLDDT> (the average pLDDT value along the sequence) since both pTM and NOESY recall are global rather than per-residues scores. The Bayesian probability of a model given NOE data is then:

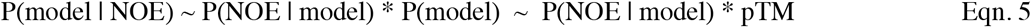

For scoring models, we use:

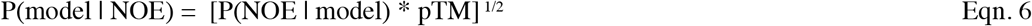

This metric incorporates both the agreement of the model with NOE data and the model’s reliability as estimated by pTM.

Similarly, to estimate P(model | RCI) for each model, we calculate the SCC between per-residue pLDDT scores and RCI (chemical shift) values:

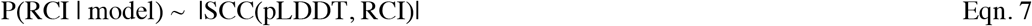

This score evaluates the agreement between pLDDT, a per-residue reliability estimate from AF2 structure models, and the per-residue NMR RCI values derived from chemical shift data, an experimental estimate of conformational flexibility. Un-normalized Bayesian likelihood is then estimated as:

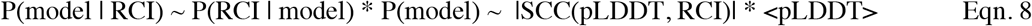

where both |SCC(pLDDT, RCI)| and <pLDDT> are scaled between 0.0 and 1.0. We used SCC instead of PCC due to the observed nonlinear relationship between pLDDT and RCI, and because SCC is less influenced by disordered tails. Vranken and co-workers report PCC(pLDDT, S^2^_RCI_) = 0.673 for a database of 76,711 residues^13^, supporting its use as a binary indicator of order vs disorder, although the AF2 pLDDT metric does not linearly correlate with the degree of dynamics within highly disordered regions of proteins. Previous studies have also shown that RCI correlates with local rigidity inferred from NMR structures based on hydrogen bond network satisfaction^76^, a correlation that was used as an estimate of NMR model accuracy.

For the purpose of scoring models vs RCI data, we use:

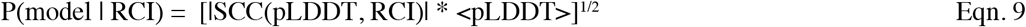

Each model is given an **AF-NMR selection score** using the Bayesian estimate of model likelihood given both RCI and NOE:

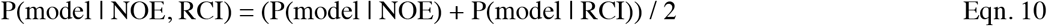

In this study, we define the **AF-NMR Bayesian model selection metric** as:

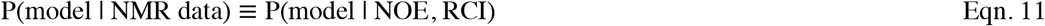

This metric enables quantitative selection of models that best match experimental data, and can also be extended to incorporate additional experimental data, such as RDCs.

### Conformational selection and state combination assessment for multi-state modeling

To identify conformer ensembles that best fit experimental data, models generated by enhanced sampling are grouped using PCA-based hierarchical clustering and scored with the AF-NMR Bayesian model selection metric. The top five models from each cluster are selected as candidate single-state ensembles. These ensembles are ranked from high to low by their average Bayesian Model Selection scores, and labeled sequentially with Greek letters (e.g., ensembles α5, β5, γ5 and clusters α, β, γ, where 5 refers to the number of conformers in the corresponding ensemble). The highest-scoring five-conformer ensemble, α5, is assigned as state 1.

To evaluate whether a multi-state ensemble better explains the data, we iteratively test combinations of the potential states. The RMSF agreement between the atomic coordinates of state 1 (RMSF_ENS_) and the RMSF estimated from RCI data (RMSF_RCI_) is first evaluated as:

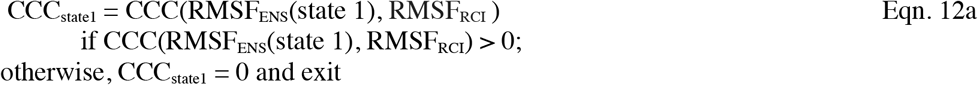

where the CCC function measures the agreement between the model-derived C*α* RMSF_ENS_ and RMSF_RCI_. Next, we assess whether adding a second candidate single-state ensemble *x* improves the correlation:

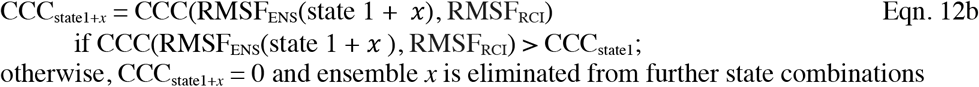

Only ensembles that improve these CCCs are retained, and the ensemble that gives the highest combined CCC becomes state 2. The process of adding and testing new ensembles is repeated until no additional states improve the fit to experimental data.

A comparison plot of RMSF_ENS_ of the final assigned state(s) vs RMSF_RCI_ is evaluated. If most differences fall within the 95% CI, then these state(s) define the solution NMR structure. If at least two distinct states are identified, the result is a multi-state structure. Models with large deviations beyond the 95% CI and a final combined CCC < 0.4 suggest an inaccurate structural representation, requiring alternative modeling strategies to generate more diverse conformers.

### Molecular graphics, secondary structure analysis, and pocket analysis

Structural visualization and molecular graphics were done using *ChimeraX*^*77*^ and *PyMol*^78^. The *Define Secondary Structure of Proteins (DSSP)*^*79*^ server (https://2struc.cryst.bbk.ac.uk/twostruc) was used for identifying regular secondary structures. The *CastpFold* server^52^ was used for surface pocket analysis.

## Abbreviations

AF2: AlphaFold2 multimer
AFsample: AlphaFold2 with enhanced sampling
AI: Artificial Intelligence
CCC: Lin’s Concordance Correlation Coefficient
CDK2AP1: Cyclin-Dependent Kinase 2-Associated Protein 1
CI: statistical Confidence Interval
LDDT: Local Difference Distance Test score, a superimposition independent metric of structural similarity
NOE: Nuclear Overhauser Effect
GLuc: *Gaussia* luciferase
NOESY: NOE SpectroscopY
pLDDT: predicted per-residue LDDT from AF2
<pLDDT>: pLDDT score averaged over all residues in the protein sequence
pLDDT_avg_: average per-residue pLDDT across a set of models
PSVS: Protein Structure Validation Software suite
pTM: predicted TM score for overall protein structure from AF2
RCI: Random Coil Index for assessing conformational flexibility from chemical shift data
RCI_0.6_: Random Coil Index scaled by 1/0.6
RDC: Residual Dipolar Coupling
RMSD: Root Mean Squared Deviation
RMSF: Root-Mean-Squared Fluctuation of atomic positions
RMSF_ENS_: RMSF for an ensemble of conformers
RMSF_RCI_: Random Coil Index scaled for RMSF comparison
RPF-DP score: Recall, Precision, F-measure, and Discrimination Power score
SA: surface area
SCC: Spearman’s correlation coefficient
SEM: standard error of the mean
TM score: template model score, a measure of structural similarity.

## Acknowledgments

We thank T. Benavides, N. Dube, A. Ertekin, K. Fraga, Y. Kuroda, E. Li, B. A. Shurina, G.V.T. Swapna, R. Tejero, and N. Wu for helpful discussions, and S. Collen for his efforts in installing and maintaining AFsample on the NPL CPU / GPU cluster of the RPI Center for Computational Innovations. This research was supported by grant R35-GM141818 (to GTM) from the National Institutes of Health, National Institute of General Medical Sciences.

## Author Contributions

YJH, TAR, LES, and GTM conceptualized the study and analyzed data. TAR, LES, and NK provided experimental data. NK provided expertise with GLuc NMR data. YJH developed computer codes. All authors contributed in writing and editing the manuscript.

## Data Availability

Atomic models of GLuc State 1 and State 2 (PDB_ID 9A8V, linked to data including NOESY peak lists and chemical shifts at https://zenodo.org/records/13831427) and CDK2AP1 State1 and State 2 (PDB_ID 9A8Z, linked to data including NOESY peak lists and chemical shifts at BMRB 16808) have been deposited in the Protein Data Bank. Additional data such as AFsample models and their scores are available at GitHub (https://github.rpi.edu/RPIBioinformatics/AlphaFold-NMR)

## Code Availability

Scripts for AF-NMR analysis presented here are available at GitHub (https://github.rpi.edu/RPIBioinformatics/AlphaFold-NMR)

