## Supplementary Material for "Hidden Structural States of Proteins Revealed by Conformer Selection with AlphaFold-NMR"

**Supplementary Figures and Tables**

Fig. S1. Comparison of two GLuc restraint-based NMR structures
Fig. S2. Flow chart of AF-NMR conformer selection protocol
Fig. S3. PCA of GLuc AFsample models
Fig. S4. Top models from GLuc AFsample clusters α, β, γ and δ
Fig. S5. Agreements of $RMSF_{RCI}$ and model atomic RMSFs for single- and multi-state ensembles
Fig. S6. Comparison of pLDDT and RCI data for GLuc.
Fig. S7. NOESY Double Recall Analysis of GLuc for $NMR_{7d2o}$ vs AF-NMR models.
Fig. S8. Top models from CDK2AP1 AFsample clusters α, β, γ and δ
Fig. S9. PCA of CDK2AP1 AFsample models
Fig. S10. Agreements of $RMSF_{RCI}$ and model atomic RMSFs for single- and multi-state ensembles for CDK2AP1
Fig. S11. Comparison of per-residue model uncertainty (pLDDT) with experimental chemical shift RCI data of CDK2AP1
Fig. S12. NOESY Double Recall analysis of AF-NMR multi-state structures of CDK2AP1
Fig S13. Peak doubling for HNs of Thr109 and Glu110 in CDK2AP1
Fig. S14. Pocket identification in CDK2AP1

Table S1. GLuc Double Recall analysis of AF-NMR state 1 and state 2 ensembles and $NMR_{7d2o}$
Table S2. Summary of GLuc structure quality factors for AF-NMR State 1
Table S3. Summary of GLuc structure quality factors for AF-NMR State 2
Table S4. Summary of GLuc structure quality factors for conventional NMR structure PDB ID 7d2o
Table S5. Summary of GLuc structure quality factors for conventional NMR structure PDB ID 9fla
Table S6. GLuc structural quality and NOESY RPF statistics for conformer-selected and restraint-based NMR models
Table S7. CDK2AP1 Double Recall analysis of AF-NMR state 1 and state 2 ensembles and $NMR_{2kw6}$
Table S8. Summary of CDK2AP1 structure quality factors for AF-NMR State 1-inter sulfur-π
Table S9. Summary of CDK2AP1 structure quality factors for AF-NMR State 2-intra sulfur-π
Table S10. Summary of CDK2AP1 structure quality factors for conventional NMR structure (PDB 2kw6)
Table S11. CDKAP1 structural quality and NOESY RPF statistics for conformer-selected and restraint-based NMR models

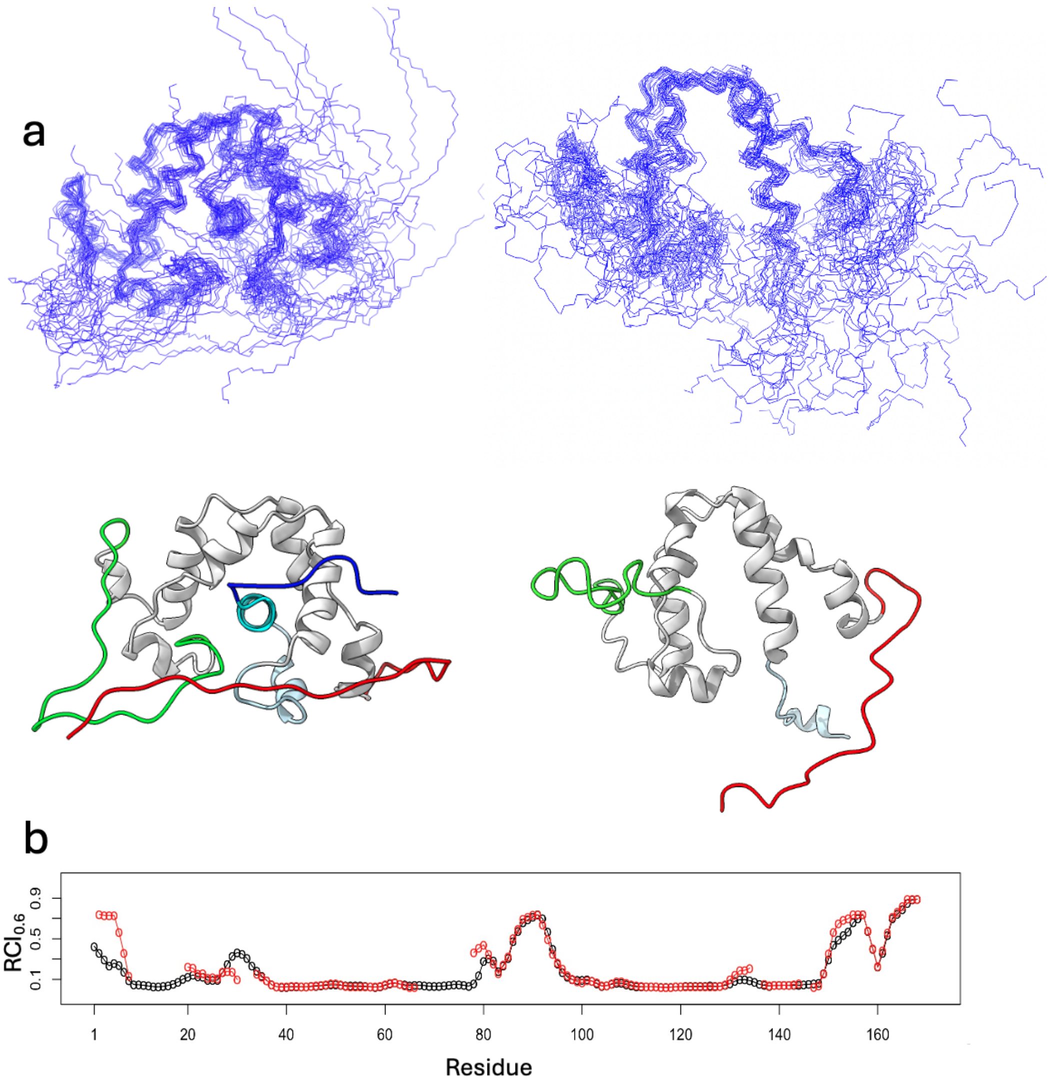


**Fig. S1. Comparison of two GLuc restraint-based NMR structures. (a)** Structures of GLuc determined by NMR from PDB IDs 7d2o [1] ($NMR_{7d2o}$, left) and 9fla [2] ($NMR_{9fla}$, right). Segment colors as defined in main text **Fig. 1a**. **(b)** $RCI_{0.6}$ plots comparing backbone flexibility for $NMR_{7d2o}$ (black, BMRB ID: 36385) and $NMR_{9fla}$ (red, BMRB ID: 34918). $NMR_{7d2o}$ has more complete backbone resonance assignments than $NMR_{9fla}$. However, for residues assigned in both BMRB entries, the resonance assignments and RCI values are highly similar.

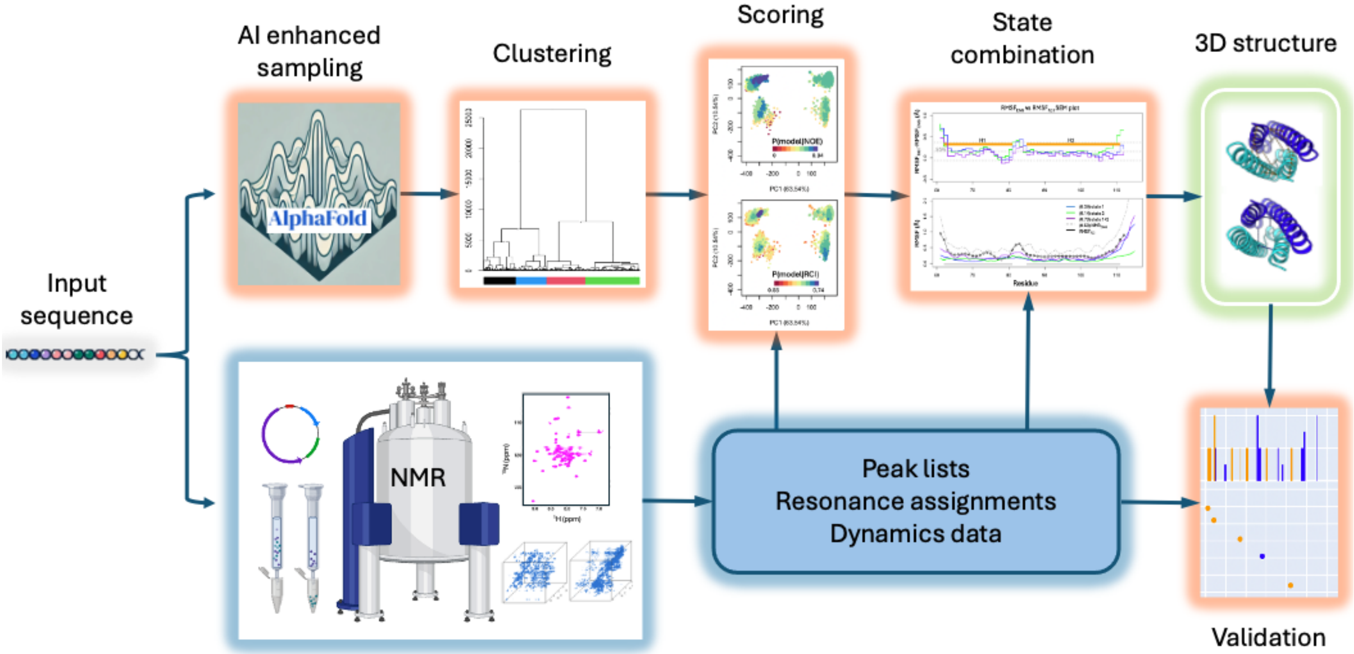


**Fig. S2**. **Flow chart of AF-NMR conformer selection protocol**. The experimental branch (blue) involves obtaining resonance assignments, NOESY peak lists, and dynamic data sensitive to multiple conformational states, including chemical shift RCI values, chemical exchange, and/or $^{15}N$ nuclear relaxation data. The computational branch (orange) employs one or more enhanced sampling methods using AI-based modeling approaches, such as AFsample[3]. The resulting models are clustered and scored against experimental data using the Bayesian model selection metric P(model | NMR data) (**Eqn. 11**) to select candidate single-state ensembles and the best scoring state 1. Dynamics data is then used to evaluate multi-state combinations (Eqn. 12). Based on fitting to these experimental data, the selected conformations are interpreted as either single-state or multi-state models of the protein structure. Multi-state models undergo additional cross-validation with NOESY Double Recall analysis. If the models do not sufficiently match the NMR data, additional enhanced sampling methods are tested, and the conformer selection and state combination process is repeated. If none of the tested enhanced sampling approaches yield models that adequately agree with the experimental data, the process is terminated.

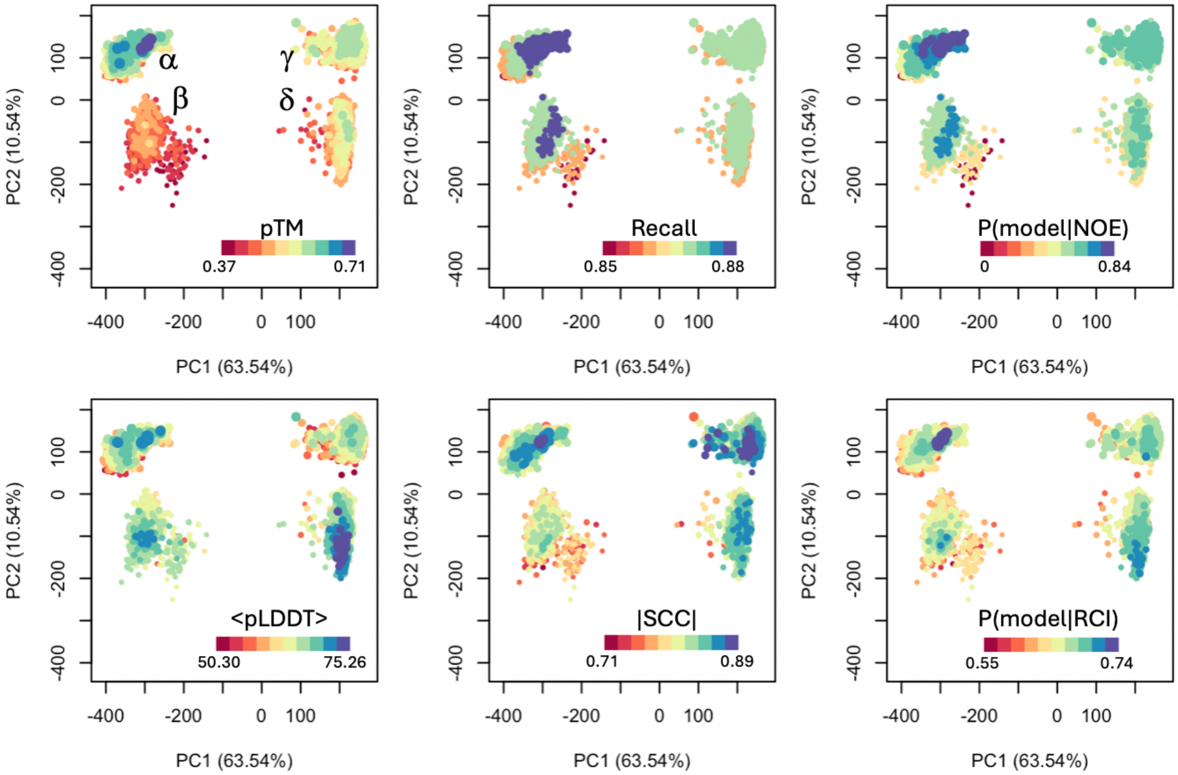


**Fig. S3. PCA of GLuc AFsample models.** 2D projections of principal component analysis (PCA) based on the C$\alpha$ - C$\alpha$ distance matrix, showing PC1 and PC2 components and the normalized contribution of each (in parentheses). Models are colored by the scores indicated in each plot (**Eqns. 4-9**). In each panel, point size corresponds to the model's pTM score, with larger points representing higher pTM values. The four PCA clusters (α, β, γ, and δ) are labeled in the top-left panel. In the pLDDT-colored plot, <pLDDT> represents the average pLDDT across all residues in each model.

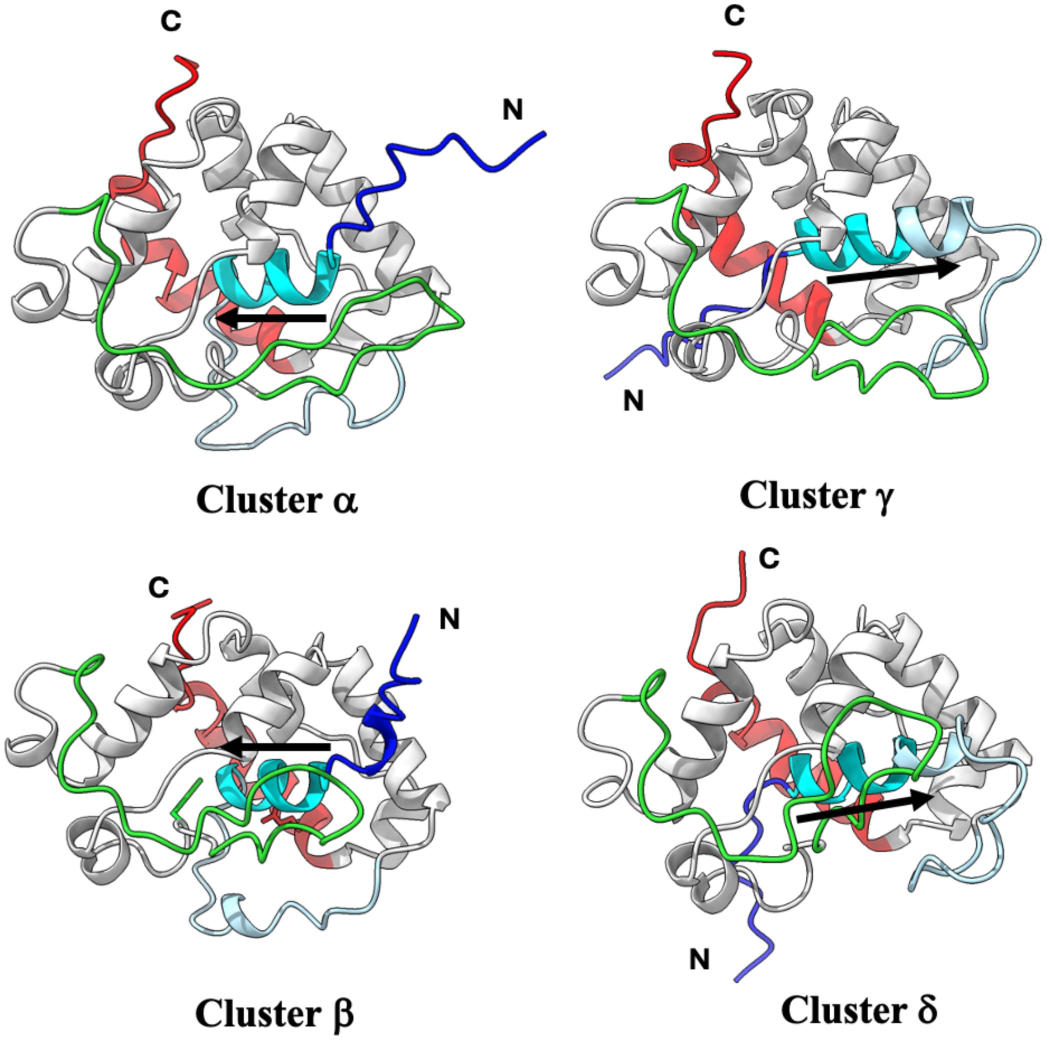


**Fig. S4. Top models from GLuc AFsample clusters (α, β, γ and δ).** Ribbon diagrams for the top-scoring model from each cluster are shown. Segment colors as defined in main text **Fig. 1a**. Clusters that separate along PC1 (e.g., α vs γ and β vs δ) primarily differ in the orientation of helix H1 relative to the core helices, as indicated by black arrows. Clusters that separate along PC2 (e.g., α vs β and γ vs δ) mainly differ in the positioning of the flexible H5/H6 thumb-shaped loop (green).

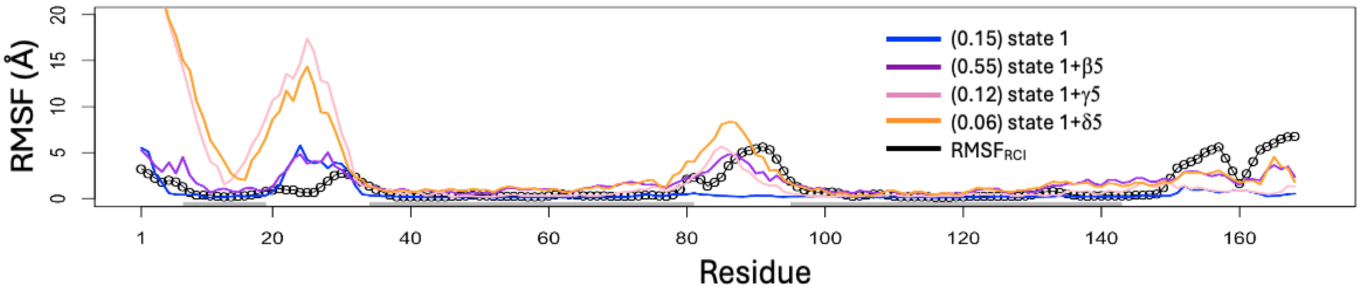


**Fig. S5. Agreement between $RMSF_{RCI}$ and C$\alpha$ atomic RMSFs for single- and multi-state ensembles.** The RCI-based RMSF ($RMSF_{RCI}$) (**Eqn. 2**), derived from chemical shifts (black circled line), is compared to the standard coordinated-based RMSF from the selected ensembles ($RMSF_{ENS}$). The "well defined" regions of $NMR_{7d2o}$ are indicated by thin gray line bars at the bottom and were used for structural alignments in $RMSF_{ENS}$ calculations. $RMSF_{ENS}$ values are shown for the ensemble of five selected models of state 1 ($\alpha$5), as well as for combined state ensembles (state 1 + $\beta$5, state 1 + $\gamma$5, and state 1 + $\delta$5). Unfiltered original CCC values, indicating the agreement between $RMSF_{ENS}$ and $RMSF_{RCI}$, are shown in parentheses.

***Conformer selection and identification of states of GLuc.*** The models generated by AFsample were evaluated against experimental NMR data using the Bayesian Model Selection metric, P(model | NOE, RCI) ( **Eqn. 10** in Methods). The top 5 highest-scoring models in each PCA cluster were selected to represent the candidate single-state ensembles. They are ordered and then named based on their average Bayesian Model Selection scores: α5 (0.770-0.787), β5 (0.690-0.697), γ5 (0.676-0.682), and δ5 (0.660-0.664).

Following the conformational selection protocol outlined in Methods, ensemble α5, with the highest Bayesian model selection score, was designated state 1. Structural variability, measured by root-mean-squared C$\alpha$ fluctuations ($RMSF_{ENS}$), was compared against experimental RCI-based RMSF ($RMSF_{RCI}$) using **Eqn. 12a**, yielding $CCC_{state1} = 0.15$. To assess whether a two-state model has better agreement, state 1 was combined with each of the other ensembles, and CCC scores for combined states were calculated (**Eqn. 12b**):

$CCC(RMSF_{ENS}(\text{state 1} + \beta5), RMSF_{RCI}) = 0.55$,

$CCC(RMSF_{ENS}(\text{state 1} + \gamma5), RMSF_{RCI}) = 0.12$,

$CCC(RMSF_{ENS}(\text{state 1} + \delta5), RMSF_{RCI}) = 0.06$.

$RMSF_{ENS}$ vs sequence profiles for each state combination are shown in **Supplementary Fig. S5**. Compared to state 1 alone, state 1 + β5 significantly improved agreement with the RCI-derived flexibility, $RMSF_{RCI}$, whereas adding γ5 or δ5 worsened the agreement. Combinations with γ5 and δ5, characterized by flipped H1 orientations (**Supplementary Figure S4**), had the largest RMSF differences, particularly in helix H1 (**Supplementary Fig. S5**), a region experimentally determined to be relatively rigid based on both RCI and $^{15}$N-$^{1}$H NOE data (**Figure 1c,d**). Additionally, NOESY Double Recall analysis did not support the flipped H1 orientation of γ5 and δ5, further indicating that these clusters do not correspond to significantly populated states. Based on these results, ensemble β5 was assigned as state 2, the second significantly populated conformational state, with little or no contributions of clusters $\gamma$ and $\delta$ under the conditions of the NMR measurements. The final AF-NMR ensemble, consisting of the ten combined models from state 1 and state 2, provides the best agreement with the experimental NOESY and RCI data.

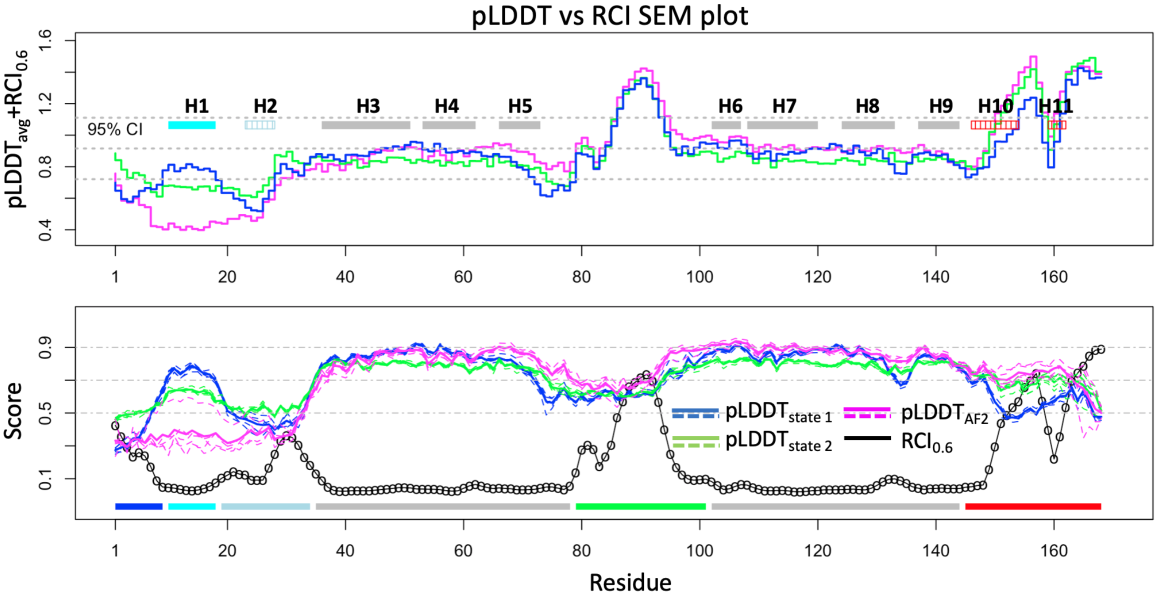


**Fig. S6. Comparison of pLDDT and RCI data for GLuc.** (Top) Comparison SEM plot of pLDDT vs RCI (see Methods), shown as the differences of $pLDDT_{avg}$ - $RCI_{0.6}$. The positions of nine helices (H1 - H9) identified in the solution NMR structure[1] and two additional helices identified by AF2 (H10 and H11) are marked with rectangles. The three most flexible helices H2, H10 and H11 are indicated with striped lines. The gray dashed lines represent the mean and 95% confidence interval (CI) (**Eqn. 3)**. (Bottom) Per-residue pLDDT and $RCI_{0.6}$ values plotted along the sequence. Dashed lines represent individual models, while the solid line denotes the average per-residue $pLDDT_{avg}$. Reference pLDDT values of 0.5, 0.7, and 0.9 are shown as gray dot-dashed lines. The color scheme from **Fig. 1a** is applied along the bottom for residue segments. Spearman correlation coefficients (SCCs) between RCI and $pLDDT_{avg}$ are -0.75 (-0.77 to -0.72) for AF-NMR state 1 and -0.67 (-0.71 to -0.64) for state 2, both showing significantly improved correlation compared to standard AF2 (-0.56, -0.59 to -0.51), mainly due to higher pLDDTs from the corrected packing of H1. This score, SCC($pLDDT_{avg}$, RCI), is described in the Methods.

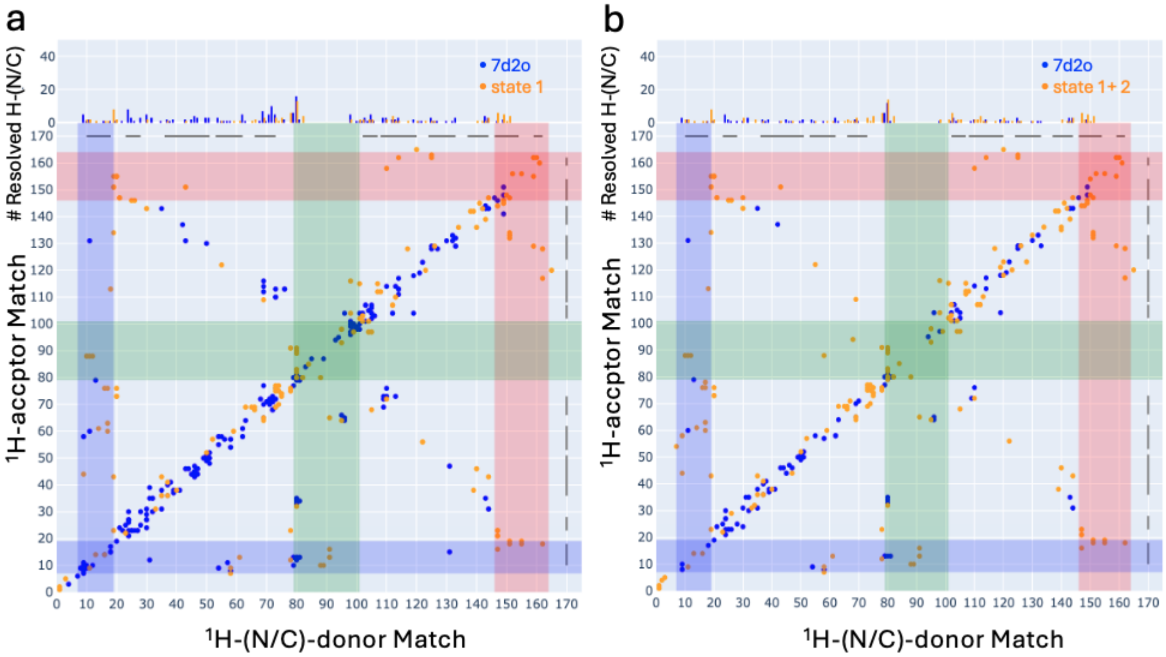


**Fig. S7. NOESY Double Recall Analysis of GLuc for $NMR_{7d2o}$ vs AF-NMR models. (a)** Double Recall plot comparing $NMR_{7d2o}$ vs AF-NMR state 1 and **(b)** $NMR_{7d2o}$ vs AF-NMR state 1 + 2 combined. The 2D contact maps plot inter residues NOEs from 3D $^{15}N/^{13}C$-resolved NOESY spectra, showing interactions uniquely explained by $NMR_{7d2o}$ (blue), AF-NMR models (orange), or both (brown). The top 1D histogram displays the per-residue count of uniquely explained NOEs, with resolved donor-matched peaks from ensemble A-only (blue) and ensemble B-only (orange). Scaling is consistent with **Fig. 1b**. Key structural interactions are highlighted by stripes: helix H1–core contacts (blue), helix H10/H11 to core contacts (red), and interactions involving the H5/H6 loop (green). The combined state 1 + 2 ensemble explains a greater number of NOEs than the single-state $NMR_{7d2o}$ model (indicated by orange dots in panel b), particularly long-range contacts between H1 (residues 7-19), H10/H11 (residues 146-164), and core residues, as well as interactions involving the thumb-shaped H5/H6 loop.

***Supplemental NOESY Double Recall Analysis for GLuc***.

We used NOESY Double Recall analysis to compare the conventional $NMR_{7d2o}$ ensemble with AF-NMR models to assess whether the selected conformational states better explain the NOESY data. (**Fig. S7, Supplementary Table S1**).

**Comparison of $NMR_{7d2o}$ v. state 1 (Fig. S7a).** AF-NMR state 1 accounts for 185 NOEs (101 long-range NOEs) that were not explained by $NMR_{7d2o}$, supporting state 1 as a significantly populated conformational state under NMR conditions (**Supplementary Table S1**). However, there are 70 long-range NOEs consistent with $NMR_{7d2o}$, but not consistent with AF-NMR state 1, indicating that state 1 does not fully capture all NOE-supported interactions. Key structural insights:

- Helix H1 interactions (blue stripe): Many NOEs exclusive to state 1 involve helix H1 (residues 7–19) interacting with the protein core, which are not accounted for by standard AF2 models (**cf. Fig. 1b,** of main text).
- C-terminal helices (red stripe): Additional NOEs between H10/H11 and core helices H1, H2, H3, H7, and H8 were identified in state 1; these are not accounted for by the $NMR_{7d2o}$ model.

- H5/H6 loop interactions (Green stripe): Seven NOEs between Tyr80 and Gly90 define the thumb-shaped feature of the H5/H6 loop,  which was not previously characterized in $NMR_{7d2o}$.

Compared to standard AF2 models (cf. **Fig. 1b,** main text), state 1 explains 266 more NOEs, confirming improved agreement with experimental NOESY data.

**Comparison of $NMR_{7d2o}$ vs AF-NMR state 1 + state 2 (Fig. S7b).** Adding state 2 further improves agreement with the NOESY data, capturing additional interactions missing in state 1 alone. The combined state 1 + state 2 ensemble explains 147 NOEs (37 long-range NOEs) that were uniquely assigned to $NMR_{7d2o}$ when only state 1 was considered. The reduction in blue dots indicates that state 2 captures NOEs missing in state 1, further improving the fit to experimental data. Newly explained interactions:

- H5/H6 loop (Green stripe): Additional interactions between the thumb-shaped loop and helices H1 and H7 are captured by state 2.
- Core helix interactions: State 2 reduces anti-diagonal blue dots, meaning it explains more NOEs assigned to long-range contacts within the core structure that were missing in state 1 alone.

The final comparison across models confirms that 116 NOEs (long-range) are uniquely explained by the combined AF-NMR state 1 + state 2 ensemble, while only 33 NOEs (long-range) remain exclusive to $NMR_{7d2o}$. Many of these 33 NOEs involve hydrogen atoms of Thr79, Tyr80, and Glu81 interacting with Ala31, Lys33, Lys34, and Leu35, suggesting a possible subpopulation of additional "open state 2" conformers in which the H5/H6 loop interacts with helix H2.

In summary, NOESY Double Recall analysis confirms that the AF-NMR state 1 + state 2 ensemble explains the experimental NOESY data better than the conventional $NMR_{7d2o}$ structure. The combined ensemble captures key structural features that were missed in single-state models, supporting the utility of AF-NMR conformational selection for resolving multiple conformational states. However, some unexplained NOEs suggest the presence of other minor conformational states in dynamic equilibrium, which may require further enhanced sampling or additional experimental validation.

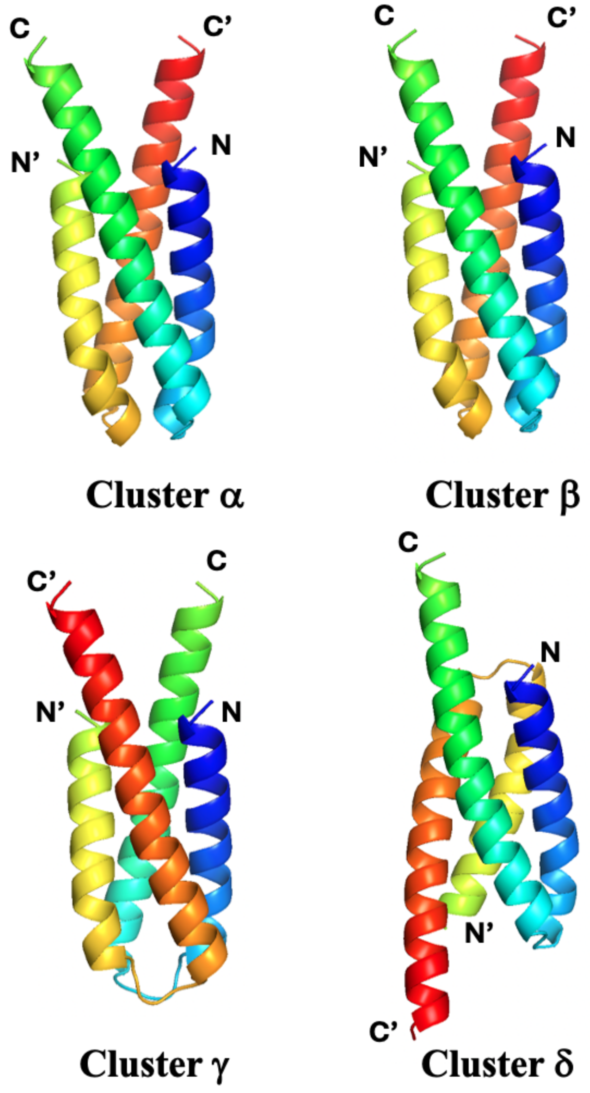


**Fig. S8. Top models from CDK2AP1 AFsample clusters.** Clusters α and β form right-handed head-to head four-helix bundles, while cluster γ adopts a left-handed head-to-head four-helix bundle. Cluster δ consists of right-handed head-to-tail four-helix bundles. The $^{13}C/^{15}N$ edited- and X-filtered NOESY data do not support models from clusters γ and δ. Chain A is shown in blue to green, and Chain B in yellow to red. Helix 1 (H1) of Chain A from each cluster is aligned for comparison.

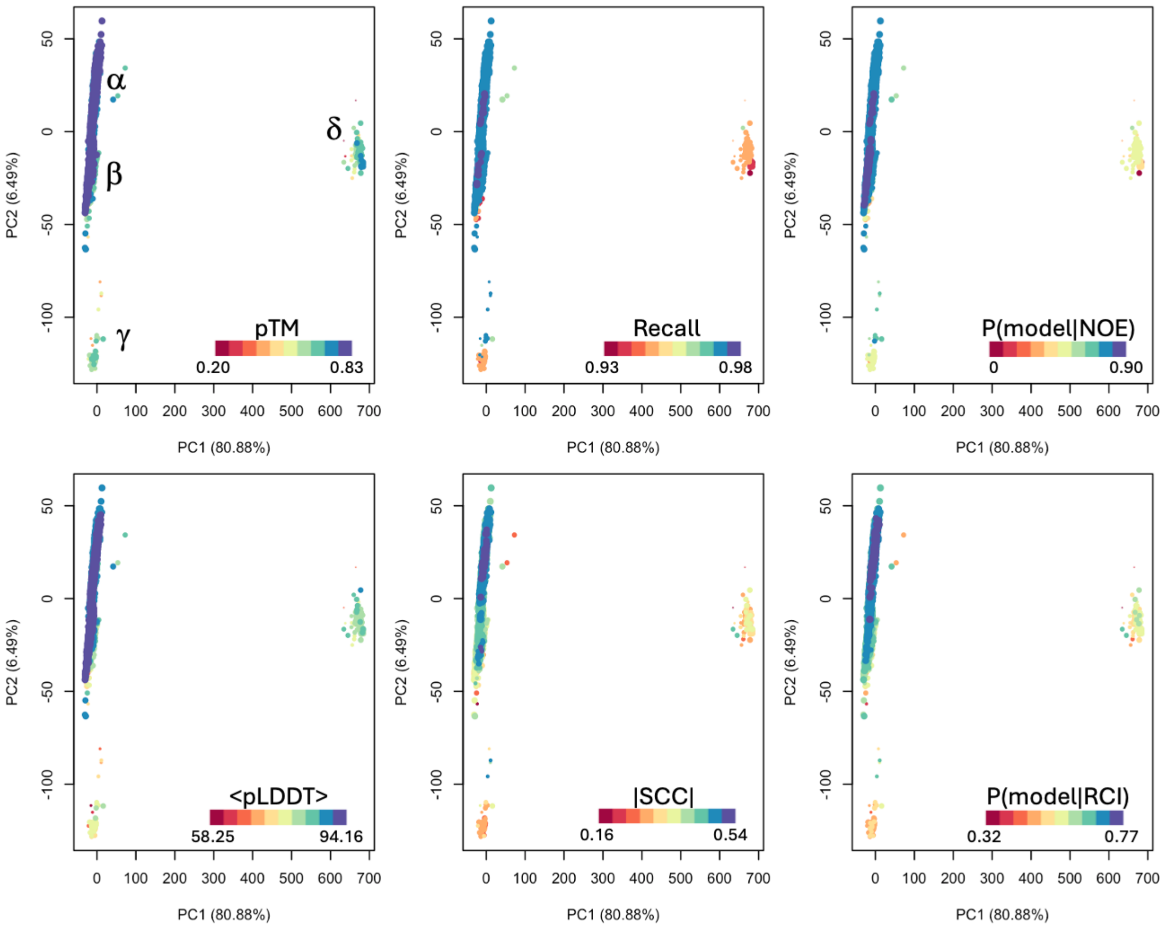


**Fig. S9. PCA of CDK2AP1 AFsample models.** Two-dimensional projections of principal component analysis (PCA) based on the C$\alpha$- C$\alpha$ distance matrix. Models are colored according to the scores indicated in each plot (**Eqn. 4-9**). The size of each point represents the pTM score, with larger points corresponding to higher pTM values. The four PCA clusters are labeled $\alpha$, $\beta$, $\gamma$, and $\delta$ in the top left panel.

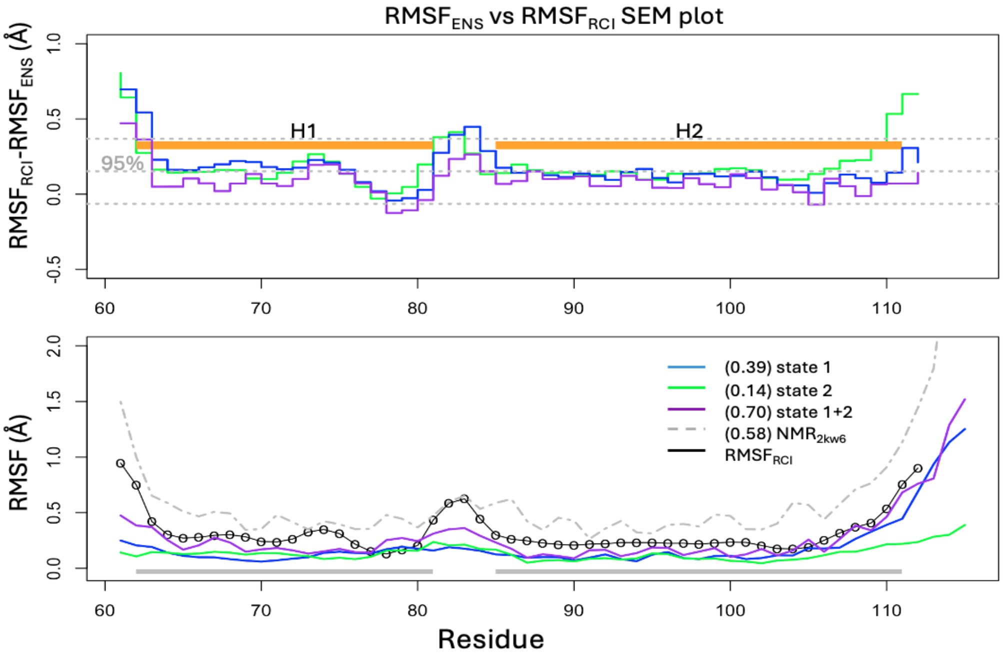


**Fig. S10. Agreements of $RMSF_{RCI}$ and model atomic RMSFs for single- and multi-state ensembles for CDK2AP1.** (Top) $RMSF_{ENS}$ vs $RMSF_{RCI}$ (**Eqn. 2)** comparison SEM plots for state 1, state 2 and the combined state 1+2 ensemble. The two helices are shown as orange line bars. (bottom) Per-residue $RMSF_{ENS}$ values for state 1, state 2, the combined state 1+2, and $NMR_{2kw6}$, compared with the $RMSF_{RCI}$. The regions used for structural alignment in RMSF calculations are shown as gray line bars at the bottom. CCCs between $RMSF_{ENS}$ for each ensemble and $RMSF_{RCI}$ are indicated in parentheses.

***Conformer selection and identification of states of CDK2AP1.*** The top 5 highest-scoring models in each PCA cluster were identified using the Bayesian Model Selection metric. These candidate single-state ensembles were ranked from high to low based on their average Bayesian Model Selection scores and subsequently named from α to δ: α5 (0.767-0.774), β5 (0.745-0.750 ), γ5 (0.590-0.624), δ5 (0.545-0.559). The inter-chain X-filtered NOESY data only supports the right-handed head-to-head four-helix bundle packing observed in the α and β clusters, whereas clusters γ and δ were eliminated due to their inconsistency with the NOESY data. However, this does not rule out the possibility that clusters γ and δ are lowly populated states that could only be observed under different experimental conditions or with alternate experimental measurements.

Following the protocol outlined in the Methods for identifying and combining conformational states to best fit the NMR data, we first assigned the highest scoring ensemble (α5) as state 1. The agreement between NMR-derived $RMSF_{RCI}$ data and the $RMSF_{ENS}$ for state 1 was evaluated using **Eqn. 12a**, yielding $CCC(RMSF_{ENS}(\text{state 1}), RMSF_{RCI}) = 0.39$, indicating moderate agreement. To assess whether a multi-state model would better explain the data, we calculated the CCC for a combination of state 1 of with ensemble $\beta 5$ using **Eqn. 12b**, yielding $CCC(RMSF_{ENS}(\text{state 1} + \beta 5), RMSF_{RCI}) = 0.70$, demonstrating a significant improvement in agreement compared to state 1 alone. Accordingly, β5 was assigned as state 2, the second significantly-populated AF-NMR conformational state. The improved CCC score confirms that a two-state ensemble provides a better fit to the NMR data than any single-state model.

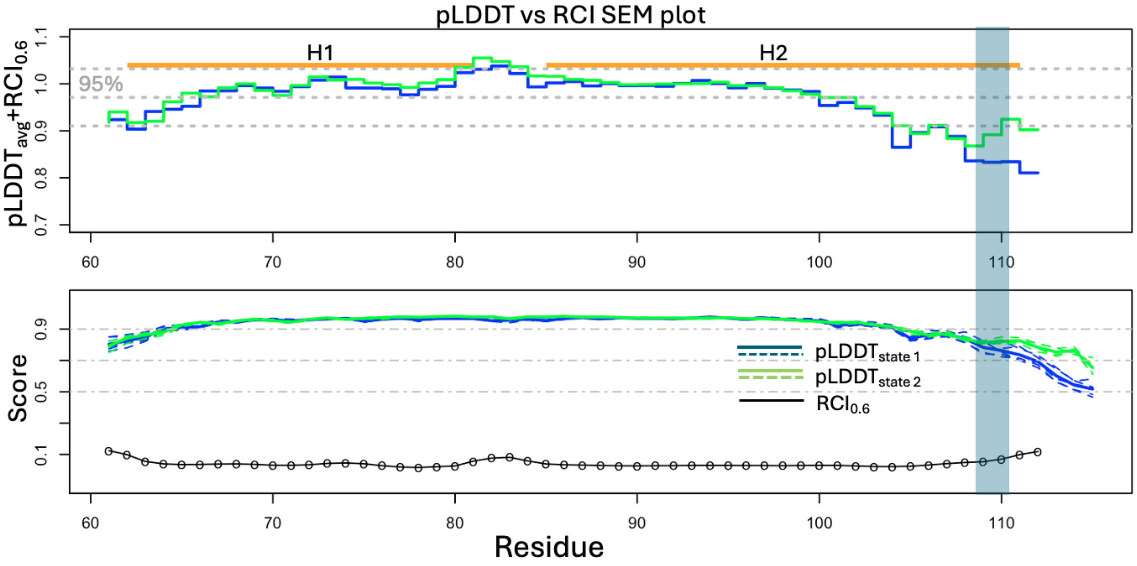


**Fig. S11. Comparison of pLDDT with RCI data for CDK2AP1.** (Top) pLDDT vs RCI comparison SEM plot (see Methods) showing the difference between $RCI_{0.6}$ and $pLDDT_{avg}$. The locations of two helices are indicated as orange line bars, while the mean and 95% confidence interval (CI) (**Eqn. 3)** are shown as gray dashed lines. (Bottom) Per-residue pLDDT (dashed line for individual models, solid line for the average per-residue pLDDT, $pLDDT_{avg}$) and $RCI_{0.6}$ scores along the sequence. Reference pLDDT values of 0.5, 0.7, 0.9 are shown as gray dot dashed lines. Spearman correlation coefficients (SCCs) between RCI and $pLDDT_{avg}$ are -0.48 (-0.48 to -0.45) for AF-NMR state 1 and -0.40 (-0.43 to -0.38) for state 2. Positions of residues Thr109 and Glu110, which exhibit HN peak doubling, are highlighted in light blue.

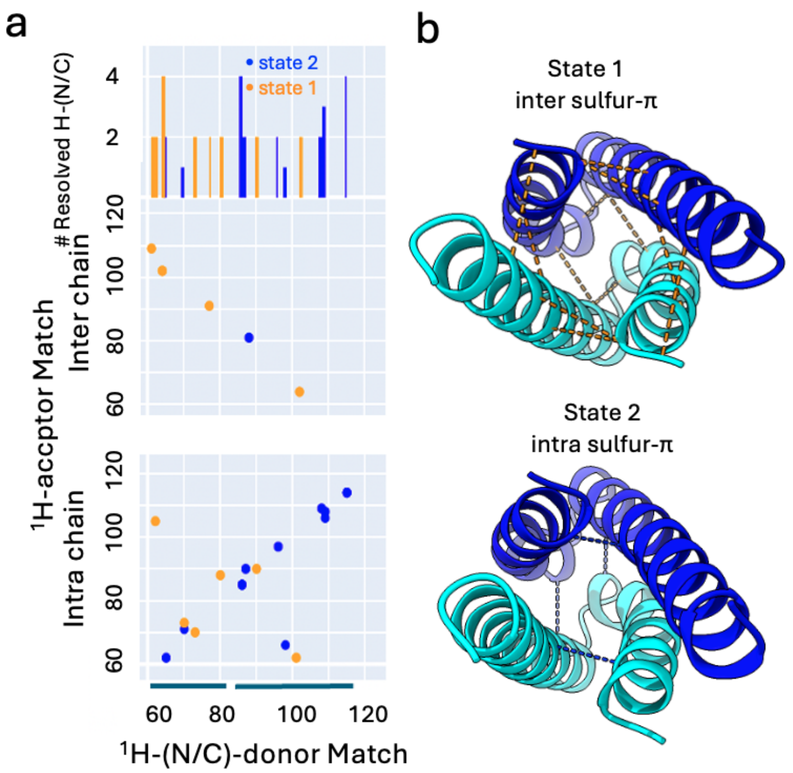


**Fig. S12. NOESY Double Recall analysis of AF-NMR multi-state structures of CDK2AP1.** (**a**) Double Recall plot of AF-NMR state 1 vs state 2, illustrating key interresidue contacts that distinguish these states; orange dots support the inter sulfur-pi state 1, blue dots support the intra sulfur-pi state 2. The "Inter chain" section maps NOE contacts between chains, while the "Intra chain" section maps NOE contacts within a single chain. NOEs with short interproton distances from both chains are counted. (Top) The 1D histogram shows the per-residues count of resolved H-(N/C) NOEs, meaning those for NOESY peaks which are not overlapped in the $^{1}$H-(X) dimension of the X-resolved NOESY spectrum, where X is $^{15}$N or $^{13}$C, except for equivalent inter chain NOEs. The locations of the two helices are shown as two line bars at the bottom. (**b-top**) Structure of state 1 long-range contacts marked by orange dashed lines; corresponding to the 8 long-range NOEs consistent with state 1 but inconsistent with state 2 (orange dots in panel **a**) with the mapping ratio: 1 dot -> 2 lines. (**b-bottom**) Structure of state 2 with long-range NOEs indicated as blue dashed lines, corresponding to 3 long range NOEs consistent with state 2 but inconsistent with state 1 (blue dots in panel **a**). The dashed lines connect the Cα-Cα atoms of the corresponding residues with the state-specific NOEs. Long range contacts unique to state 1 (orange): inter-chain: 61-109 (inter-chain), 62-101 (intra-chain), 62-105 (intra-chain), 64-102 (inter-chain), 77-91 (inter-chain), and 80-88 (intra-chain). Long range contacts unique to state 2 (blue): 66-98 (intra-chain) and 81-88 (two inter-chain NOEs).

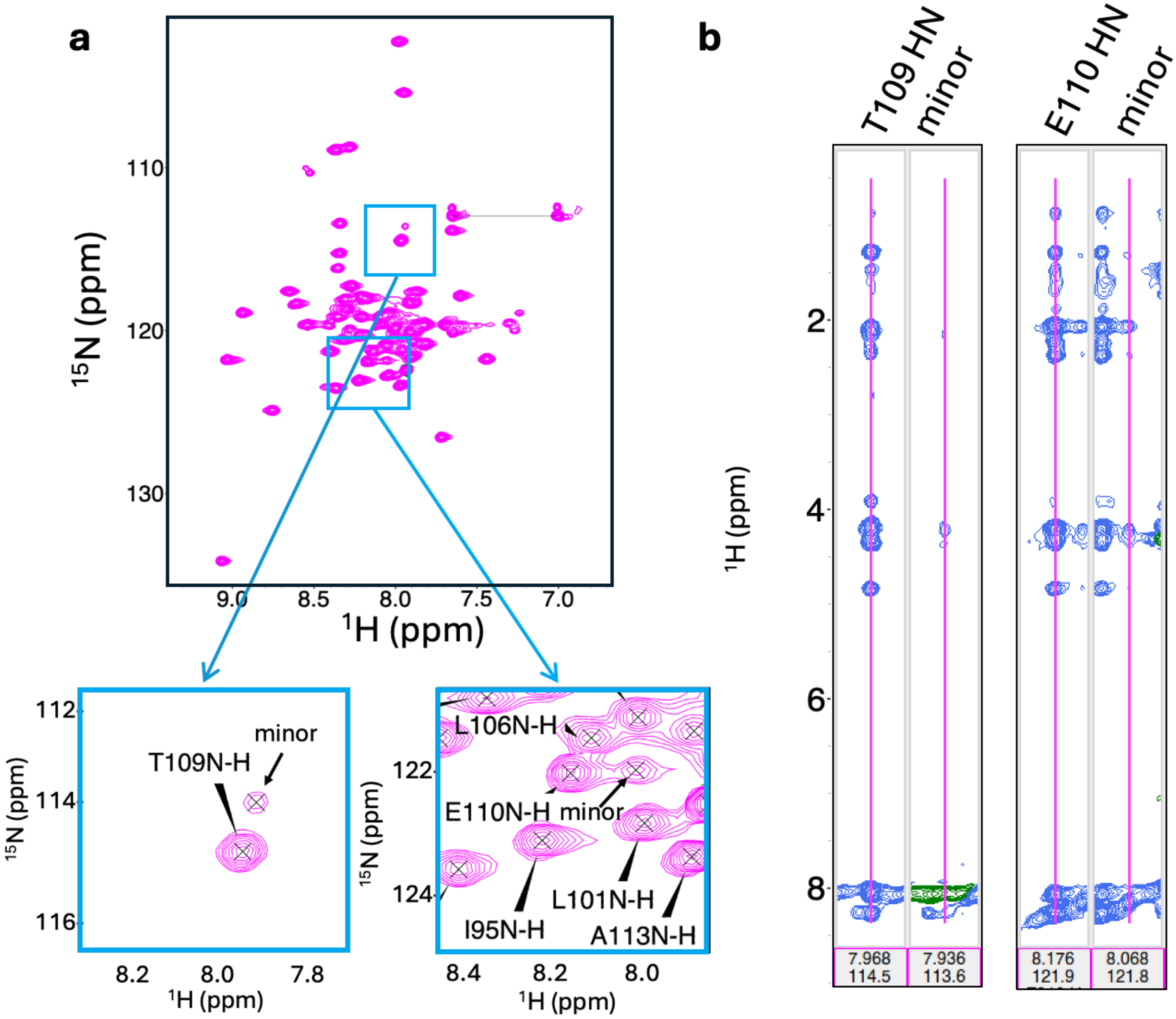


**Fig. S13. Spectral peak doubling for HNs of Thr109 and Glu110 in CDK2AP1. (a)** $^{1}$H-$^{15}$N SOFAST-HMQC spectra showing selected regions where Thr109 and Glu110 HN peaks appear as major and minor populations. **(b)** CN-NOESY spectra (mixing time = 120 ms) showing selected $^{15}$N-NOESY strips for Thr109 and Glu110 major and minor HNs, displayed as strip plots using the program POKY[4]. Assignments of the minor HN peaks were validated by their $^{1}$H-$^{15}$N edited $^{1}$H-$^{1}$H NOEs to intra-residue Hα, Hβ, and Hγ2 for Thr109, and Hα, Hβs, and Hγs for Glu110, which match those of the major population. Minor peaks have ~15-20% of the intensity of the major peaks.

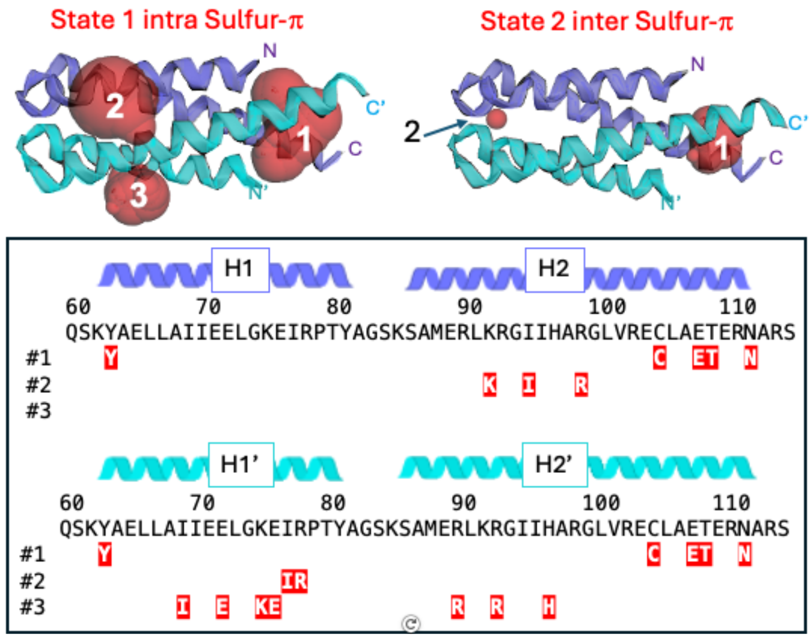


**Fig. S14. Pocket identification in CDK2AP1. (a)** Pockets identified and characterized by *CASTpFold*[5] are shown in translucent red (**Fig. 5e, top**). The homodimer has two-fold rotational symmetry which becomes slightly asymmetric in forming these pockets. An equivalent structure is formed by swapping the helix labels H1/H2 and H1'/H2'. **(b)** Sequence and labeled helical schematic with key residues indicated in red for each pocket (#1 - #3).

Pocket details across two states (**top**):

State 1 (model with the most open pockets by total surface area (SA), rank 2):

- pocket #1: volume 217 Å$^3$, SA 187 Å$^2$ – "end cavity" that includes residues from both chains.
- pocket #2: volume 82 Å$^3$, SA 94 Å$^2$ – "side pocket" located between helices H2 and H1'.
- pocket #3: volume 65 Å$^3$, SA 106 Å$^2$ – located in the "V" region between H1 and H2 of the same chain

State 2 (model with the most closed pockets by total SA, rank 1):

- pocket #1: volume 67 Å$^3$, SA 88 Å$^2$ (rank1)
- pocket #2: volume < 2 Å$^3$
- pocket #3: not detected

Pocket residues for each pocket include:

- pocket #1: Tyr63, Cys105, Glu108, Thr109, and Asn112 (from both chains)
- pocket #2: Lys92, Ile95, Arg99 (in H2) and Ile77, Arg78 (in H1')
- pocket #3: Ile69, Glu72, Lys75, Glu76 (in H1') and Arg90, Arg93, His97 (in H2')
- An equivalent structure is formed by swapping the helix labels H1/H2 and H1'/H2'.

Cavity volume and SA variability across 5 models:

State 1:

- pocket #1: volume 50–217 Å$^3$, SA 60-190 Å$^2$
- pocket #2: volume 3–83 Å$^3$, SA 10-94 Å$^2$
- pocket #3: volume 1–65 Å$^3$, SA 4-106 Å$^2$

State 2:

- pocket #1: volume 31–67 Å$^3$, SA 88-144 Å$^2$
- pocket #2: volume 0–89 Å$^3$, SA 0-86 Å$^2$
- pocket #3: volume < 1 Å$^3$, SA < 4 Å$^2$ (all cases)

**Table S1. GLuc Double Recall analysis of *AF-NMR* state 1 and state 2 ensembles and NMR$_{7d2o}$**

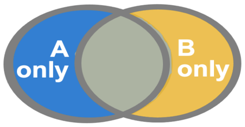


| | | # of ensemble A-only NOEs (represented by blue dots in Figs . 4a and S7) | | # of ensemble B-only NOEs (represented by orange dots in Fig. 4a and S7) | |
|---|---|---|---|---|---|
| A | B | total | long-range / resolved[a] | total | long range / resolved |
| NMR$_{7d2o}$ | state 1 | 277 | 70 / 60 | 185 | 101 / 67 |
| | state 2 | 248 | 87 / 70 | 164 | 71 / 42 |
| | **state 1+ state 2** | **128** | **33 / 27** | **218** | **116 / 77** |
| state 2 | state 1 | 175 | 55 / 40 | 158 | 89 / 58 |

[a] resolved - NOESY peaks are those that match with long-range NOEs (≥ 5 residues apart in sequence) and do not overlap in the $^{1}$H-(X) dimension of the X-resolved NOESY spectrum, where X = $^{15}$N or $^{13}$C. Unresolved NOESY peaks, by contrast, are for those long range NOEs that *do* overlap in the $^{1}$H-(X) dimension, making their assignment to a specific NOE ambiguous.

Notably, the number of both long-range and resolved NOEs increased significantly when comparing the combined state 1 + state 2 (116 & 77) to standard NMR$_{7d2o}$ model (33 & 27), highlighting a key advantage of the AF-NMR method. In conventional NMR structure determination, assigning NOESY peaks to spatial restraints in flexible regions—such as the H5/H6 loop and H11—is particularly challenging. This difficulty arises from the increased peak degeneracy in flexible regions and complications in assignments of ambiguous NOEs using iterative build-up methods in these regions. AF-NMR provided models that give a more comprehensive structural interpretation of flexible protein regions.

***Quality assessment of atomic models of GLuc's states 1 and 2***. To ensure that the AF-NMR-selected models of GLuc's two conformational states are physically reasonable, their ensembles were evaluated using knowledge-based structural quality statistics via the *Protein Structure Validation Server (PSVS)*[6]. Key validation metrics for the 5-conformer ensembles representing each state and the 19-conformer ensemble of the conventional restraint-based NMR structure are summarized in **Supplementary Table S1** above, with complete PSVS reports in **Supplementary Tables S3 – S5**. PSVS analysis reports structure quality assessment statistics as Z scores relative to values obtained for high-resolution X-ray crystal structures[6], where more values indicate better structural quality; good-quality NMR structures typically have Z scores > 0. The AF-NMR models exhibit superior knowledge-based structural quality compared to the conventional NMR$_{7d2o}$ ensemble. In particular, *ProCheck*(all) and *MolProbity* scores, which are sensitive to sidechain conformational distributions and backbone and sidechain atomic packing, respectively, are significantly better for the AF-NMR models than for the conventional NMR$_{7d2o}$ structure.

**Table S2. Summary of GLuc structure quality factors for AF-NMR State 1**

| | | | |
|---|---|---|---|
| **Number of conformers** | 5 | | |
| **RMSD Values** | | | |
| | all | Ordered[a] | Selected[b] |
| All backbone atoms | 1.0 Å | 0.3 Å | 0.2 Å |
| All heavy atoms | 1.3 Å | 0.4 Å | 0.4 Å |
| **Structure Quality Factors - overall statistics** | | | |
| | Mean score | S.D. | Z-score[c] |
| Procheck G-factor (phi / psi only) | -0.01 | N/A | 0.28 |
| Procheck G-factor [e](all dihedral angles) | 0.17 | N/A | 1.01 |
| MolProbity clashscore | 1.55 | 3.4659 | 1.26 |
| **Ramachandran Plot Summary from Procheck** | | | |
| Most favored regions | 89.7% | | |
| Additionally allowed regions | 10.3% | | |
| Generously allowed regions | 0.0% | | |
| Disallowed regions | 0.0% | | |
| **Ramachandran Plot Statistics from Richardson's lab statistics** | | | |
| Most favored regions | 96.4% | | |
| Allowed regions | 3.6% | | |
| Disallowed regions | 0% | | |

[a] Residues with sum of phi and psi order parameters >= 1.8: 5-19, 33-73, 76-165

[b] Well-defined residues used for analysis (core residues by CYRANGE ): 6..19, 34..150, 164..168

[c] With respect to mean and standard deviation for a set of 252 X-ray structures < 500 residues, of resolution <= 1.80 Å, R-factor <= 0.25 and R-free <= 0.28; a positive value indicates a 'better' score

**Table S3. Summary of GLuc structure quality factors for AF-NMR State 2**

| | | | |
|---|---|---|---|
| **Number of conformers** | 5 | | |
| **RMSD Values** | | | |
| | all | Ordered[a] | Selected[b] |
| All backbone atoms | 0.8 Å | 0.7 Å | 0.5 Å |
| All heavy atoms | 1.2 Å | 0.9 Å | 0.7 Å |
| **Structure Quality Factors - overall statistics** | | | |
| | Mean score | S.D. | Z-score[c] |
| Procheck G-factor (phi / psi only) | -0.21 | N/A | -0.51 |
| Procheck G-factor (all dihedral angles) | -0.15 | N/A | -0.89 |
| MolProbity clashscore | No clash | No clash | >3.00 |
| **Ramachandran Plot Summary from Procheck** | | | |
| Most favored regions | 84.2% | | |
| Additionally allowed regions | 15.7% | | |
| Generously allowed regions | 0.2% | | |
| Disallowed regions | 0.0% | | |
| **Ramachandran Plot Statistics from Richardson's lab statistics** | | | |
| Most favored regions | 92.4% | | |
| Allowed regions | 6.7% | | |
| Disallowed regions | 1% | | |

[a] Residues with sum of phi and psi order parameters >= 1.8: 2-20, 24-74, 76A-80, 83-85, 88-161

[b] Well-defined residues used for analysis (Core residues by CYRANGE ): 8..18, 31..163

[c] With respect to mean and standard deviation for a set of 252 X-ray structures < 500 residues, of resolution <= 1.80 Å, R-factor <= 0.25 and R-free <= 0.28; a positive value indicates a 'better' score

**Table S4. Summary of GLuc structure quality factors for conventional NMR structure PDB ID 7d2o**

| | | | |
|---|---|---|---|
| **Number of conformers** | 19 | | |
| **RMSD Values** | | | |
| | all | Ordered[a] | Selected[b] |
| All backbone atoms | 7.4 Å | 1.4 Å | 1.1 Å |
| All heavy atoms | 7.2 Å | 1.7 Å | 1.4 Å |
| **Structure Quality Factors - overall statistics** | | | |
| | Mean score | S.D. | Z-score[c] |
| Procheck G-factor (phi / psi only) | -0.24 | N/A | -0.63 |
| Procheck G-factor (all dihedral angles) | -0.63 | N/A | -3.73 |
| MolProbity clashscore | 23.66 | 1.6608 | -2.53 |
| **Ramachandran Plot Summary from Procheck** | | | |
| Most favored regions | 86.3% | | |
| Additionally allowed regions | 10.9% | | |
| Generously allowed regions | 1.2% | | |
| Disallowed regions | 1.5% | | |
| **Ramachandran Plot Statistics from Richardson's lab statistics** | | | |
| Most favored regions | 91.3% | | |
| Allowed regions | 6.6% | | |
| Disallowed regions | 2.1% | | |

[a] Residues with sum of phi and psi order parameters >= 1.8: 8-18, 23-27, 35-78, 97-134, 137-144

[b] Well-defined residues used for analysis (Core residues by CYRANGE ): 7..19, 34..81, 95..143

[c] With respect to mean and standard deviation for a set of 252 X-ray structures < 500 residues, of resolution <= 1.80 Å, R-factor <= 0.25 and R-free <= 0.28; a positive value indicates a 'better' score

**Table S5. Summary of GLuc structure quality factors for conventional NMR structure PDB ID 9fla**

| | | | |
|---|---|---|---|
| **Number of conformers** | 20 | | |
| **RMSD Values** | | | |
| | all | Ordered[a] | Selected[b] |
| All backbone atoms | 6.8 Å | 1.5 Å | 1.4 Å |
| All heavy atoms | 6.9 Å | 1.8 Å | 1.7 Å |
| **Structure Quality Factors - overall statistics** | | | |
| | Mean score | S.D. | Z-score[c] |
| Procheck G-factor (phi / psi only) | 0.12 | N/A | 0.79 |
| Procheck G-factor (all dihedral angles) | -0.04 | N/A | -0.24 |
| MolProbity clashscore | 11.83 | 1.7655 | -0.50 |
| **Ramachandran Plot Summary from Procheck** | | | |
| Most favored regions | 93.5% | | |
| Additionally allowed regions | 6.2% | | |
| Generously allowed regions | 0.3% | | |
| Disallowed regions | 0.0% | | |
| **Ramachandran Plot Statistics from Richardson's lab statistics** | | | |
| Most favored regions | 95.9% | | |
| Allowed regions | 3.8% | | |
| Disallowed regions | 0.3% | | |

[a] Residues with sum of phi and psi order parameters >= 1.8: 35-67, 97-134, 138-144

[b] Well-defined residues used for analysis (Core residues by CYRANGE ): 38..69, 95..140

[c] With respect to mean and standard deviation for a set of 252 X-ray structures < 500 residues, of resolution <= 1.80 Å, R-factor <= 0.25 and R-free <= 0.28; a positive value indicates a 'better' score

**Table S6. GLuc Structure quality and NOESY RPF statistics for conformer-selected and restraint-based NMR models**

| | ProCheck[a] (bb) | ProCheck[a] (all) | MolProbity[a] | Ramachandran Statistics[b] (percent most favored/ allowed/disallowed) | RPF[c] (Recall$_{avg}$ / Precision$_{avg}$ / F-measure$_{avg}$ / DP$_{avg}$) |
|---|---|---|---|---|---|
| state 1 | 0.28 | 1.01 | 1.26 | 96.4 / 3.6 / 0.0 | 0.88 / 0.79 / 0.82 / 0.60 |
| state 2 | -0.51 | -0.89 | > 3.00[d] | 92.4 / 6.7 / 1.0 | 0.88 / 0.79 / 0.82 / 0.60 |
| 7d2o | -0.63 | -3.73 | -2.53 | 91.3 / 6.6 / 2.1 | 0.89 / 0.79 / 0.84 / 0.67 |
| 9lfa | 0.79 | -0.24 | -0.5 | 93.5 / 6.2 / 0.3 | N/A |

[a] Z scores from the PSVS server [6] using ordered residues selected by Cyrange . [b]Percent of backbone residues in percent most favored / allowed / disallowed of Richardson Ramachandran map [7] using ordered residues selected by Cyrange . [c] RPF scores [8] are calculated for each model of the ensemble and then averaged. [d]MolProbity returns NA for models with no clash violations, and the corresponding Z score is reported as > 3.00.

***NOESY RPF Analysis for GLuc.*** The two AF-NMR states were evaluated against experimental NOESY peak list data using *RPF* analysis. RPF scores[8] were computed for each model within each state and then averaged (**Supplementary Table S6 above**). The average recall (recall$_{avg}$) for NMR$_{7d2o}$ is slightly higher than that for either state 1 or state 2 alone, likely because the restraint-based NMR$_{7d2o}$ ensemble was generated by satisfaction of NOEs arising from a mixture of both states. This approach may end up forcing the satisfaction of constraints from multiple conformations into a single incorrect/distorted model. However, Double Recall analysis demonstrates that combining state 1 and state 2 explains more NOESY peaks than either state alone, capturing structural features not identified in the conventional NMR$_{7d2o}$ models.

**Table S7. CDK2AP1 Double Recall analysis of *AF-NMR* state 1 and state 2 ensembles and NMR$_{2kw6}$**

| 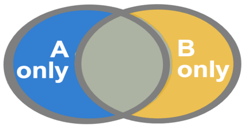  | | # of ensemble A-only NOEs | | # of ensemble B-only NOEs | |
|---|---|---|---|---|---|
| A | B | total[a] | long-range | total | long range |
| NMR$_{2kw6}$ | state 1 | 40(24) | 4(2) | 34(28) | 14 (8) |
| | state 2 | 45(20) | 19(4) | 36(29) | 10(3) |
| | **state 1+ state 2** | 23(16) | 3(2) | 42(34) | 16(8) |
| state 2 | state 1 | 23(12)[b] | 3(0) | 22(6) | 16(4) |

[a] NOEs with short interproton distances for both chains are assessed. [b] Values in parentheses are from the original NOESY peak lists before additional structure-guided NOESY peak picking.

***NOESY Double Recall analysis of CDK2AP1 for the conventional NMR$_{2kw6}$ models vs AF-NMR models.*** As a cross validation of the two-state ensemble of CDK2AP1 determined by the AF-NMR process, we also carried out NOESY Double Recall analysis for the conventional NMR$_{2kw6}$ models vs AF-NMR models (**Supplementary Table S7 above**). Only 23 NOEs (3 long range) are explained by NMR$_{2kw6}$ but not the combined state model ensemble, while 42 NOEs (16 long range) are explained by the combined state ensemble but not by the NMR$_{2kw6}$ model.

***Quality assessment of atomic models of CDK2AP1's states 1 and 2.*** The structural ensembles of state 1 and state 2 were evaluated using *PSVS*[6]. Key structure quality statistics for the 5 models of each state, along with the 20-conformer ensemble of the conventional NMR structure (2kw6), are summarized in **Supplementary Table S11**, with full PSVS reports provided in **Supplementary Tables S8 – S10**. This analysis confirms that the AF-NMR models exhibit superior knowledge-based structure quality metrics compared to the conventional restraint-based NMR models. In particular, *MolProbity* scores, which assess backbone and sidechain atomic packing, are significantly higher for the AF-NMR models than for NMR structure 2kw6.

**Table S8. Summary of CDK2AP1 structure quality factors for AF-NMR State 1 (inter sulfur-π)**

| | | | |
|---|---|---|---|
| **Number of conformers** | 5 | | |
| **RMSD Values** | | | |
| | all | | |
| All backbone atoms | 0.3 Å | | |
| All heavy atoms | 0.5 Å | | |
| **Structure Quality Factors - overall statistics** | | | |
| | Mean score | S.D. | Z-score[a] |
| Procheck G-factor (phi / psi only) | 0.77 | N/A | 3.34 |
| Procheck G-factor [e](all dihedral angles) | 0.61 | N/A | 3.61 |
| MolProbity clashscore | No clash | No clash | >3.00 |
| **Ramachandran Plot Summary from Procheck** | | | |
| Most favored regions | 100.0% | | |
| Additionally allowed regions | 0.0% | | |
| Generously allowed regions | 0.0% | | |
| Disallowed regions | 0.0% | | |
| **Ramachandran Plot Statistics from Richardson's lab statistics** | | | |
| Most favored regions | 100% | | |
| Allowed regions | 0% | | |
| Disallowed regions | 0% | | |

[a] With respect to mean and standard deviation for a set of 252 X-ray structures < 500 residues, of resolution <= 1.80 Å, R-factor <= 0.25 and R-free <= 0.28; a positive value indicates a 'better' score

**Table S9. Summary of CDK2AP1 structure quality factors for AF-NMR State 2 (intra sulfur-π)**

| | | | |
|---|---|---|---|
| **Number of conformers** | 5 | | |
| **RMSD Values** | | | |
| | all | | |
| All backbone atoms | 0.2 Å | | |
| All heavy atoms | 0.4 Å | | |
| **Structure Quality Factors - overall statistics** | | | |
| | Mean score | S.D. | Z-score[a] |
| Procheck G-factor (phi / psi only) | 0.78 | N/A | 3.38 |
| Procheck G-factor (all dihedral angles) | 0.62 | N/A | 3.67 |
| MolProbity clashscore | No clash | No clash | >3.00 |
| **Ramachandran Plot Summary from Procheck** | | | |
| Most favored regions | 100.0% | | |
| Additionally allowed regions | 0.0% | | |
| Generously allowed regions | 0.0% | | |
| Disallowed regions | 0.0% | | |
| **Ramachandran Plot Statistics from Richardson's lab statistics** | | | |
| Most favored regions | 100% | | |
| Allowed regions | 0% | | |
| Disallowed regions | 0% | | |

[a] With respect to mean and standard deviation for a set of 252 X-ray structures < 500 residues, of resolution <= 1.80 Å, R-factor <= 0.25 and R-free <= 0.28; a positive value indicates a 'better' score

**Table S10. Summary of CDK2AP1 structure quality factors for conventional NMR structure PDB ID 2kw6**

| | | | |
|---|---|---|---|
| **Number of conformers** | 20 | | |
| **RMSD Values** | | | |
| | all | | |
| All backbone atoms | 0.9 Å | | |
| All heavy atoms | 1.5 Å | | |
| **Structure Quality Factors - overall statistics** | | | |
| | Mean score | S.D. | Z-score[a] |
| Procheck G-factor (phi / psi only) | 0.46 | N/A | 2.12 |
| Procheck G-factor (all dihedral angles) | 0.35 | N/A | 2.07 |
| MolProbity clashscore | 15.57 | 2.7774 | -1.15 |
| **Ramachandran Plot Summary from Procheck** | | | |
| Most favored regions | 96.9% | | |
| Additionally allowed regions | 3.0% | | |
| Generously allowed regions | 0.1% | | |
| Disallowed regions | 0.0% | | |
| **Ramachandran Plot Statistics from Richardson's lab statistics** | | | |
| Most favored regions | 98.5% | | |
| Allowed regions | 1.3% | | |
| Disallowed regions | 0.2% | | |

[a] With respect to mean and standard deviation for a set of 252 X-ray structures < 500 residues, of resolution <= 1.80 Å, R-factor <= 0.25 and R-free <= 0.28; a positive value indicates a 'better' score

**Table S11. CDK2AP1 Structure quality and NOESY RPF statistics for conformer-selected and restraint-based NMR models**

| | ProCheck[a] (bb) | ProCheck[a] (all) | MolProbity[a] | Ramachandran Statistics[b] (percent most favored/ allowed/disallowed) | RPF[c] (Recall$_{avg}$ / Precision$_{avg}$ / F-measure$_{avg}$ / DP$_{avg}$) |
|---|---|---|---|---|---|
| state 1 | 3.34 | 3.61 | > 3.00[d] | 100 / 0.0 / 0.0 | 0.98 / 0.88 / 0.93 / 0.77 |
| state 2 | 3.38 | 3.67 | > 3.00[d] | 100 / 0.0 / 0.0 | 0.98 / 0.89 / 0.93 / 0.77 |
| 2kw6 | 3.03 | 3.13 | -1.15 | 99.9 / 0.1 / 0.0 | 0.98 / 0.88 / 0.93 / 0.75 |

[a] Z scores from the PSVS server [6] using all residues. [b]Percent of backbone residues in percent most favored / allowed / disallowed of Richardson Ramachandran map [7] using all residues. [c] RPF scores [8] are calculated for each model of the ensemble and then averaged. [d]MolProbity returns NA for models with no clash violations, and the corresponding Z score is reported as > 3.00.

**NOESY RPF Analysis for CDKAP1.** The two AF-NMR states were evaluated against experimental NOESY peak list data using *RPF* analysis (Huang *et al.*, 2005). RPF scores were computed for each model within state 1 and state 2, then averaged (**Supplementary Table S11 above**). The RPF scores for $NMR_{2kw6}$ are comparable to those for state 1 or state 2 alone.
